# NN-800s: Brain-penetrant trimeric nanobodies enable potent TNFα inhibition via TfR1-mediated transcytosis

**DOI:** 10.64898/2026.04.28.721316

**Authors:** Tao Yin, Sanjay Metkar, Metin Yesiltepe, Luciano D’Adamio

**Affiliations:** Department of Pharmacology, Physiology & Neuroscience New Jersey Medical School, Brain Health Institute, Jacqueline Krieger Klein Center in Alzheimer’s Disease and Neurodegeneration Research, Rutgers, The State University of New Jersey, 205 South Orange Ave, Newark, NJ, 0703, USA; NanoNewron LLC, Townsend Hall T27, 000 Morris Avenue, Union, NJ 07083

## Abstract

Tumor necrosis factor alpha (TNFα) is a central mediator of neuroinflammation and synaptic dysfunction in multiple central nervous system (CNS) disorders, including Alzheimer’s disease. Although clinically approved TNFα inhibitors are highly effective in peripheral inflammatory diseases, their therapeutic application in CNS disorders is severely limited by poor blood–brain barrier (BBB) penetration.

Here, we report the development of NN-800s, a class of heterotrimeric nanobody-based biologics engineered to achieve both high TNFα neutralization potency and efficient BBB transcytosis. NN-800s consist of two humanized anti-TNFα nanobodies flanking a humanized anti-transferrin receptor 1 (TfR1) nanobody that mediates receptor-dependent transport across the BBB. These constructs exhibit picomolar TNFα inhibitory activity and achieve cerebrospinal fluid (CSF)-to-serum ratios of up to ~0.4 following systemic administration.

Importantly, NN-800s do not disrupt transferrin–TfR1 interactions and do not induce hematological toxicity in vivo in humanized Tf/TfR1 rat models, supporting a favorable safety profile. The constructs are efficiently produced in CHO cells with high purity, low endotoxin levels, and strong scalability, supporting their developability as therapeutic biologics.

Together, these data establish NN-800s as a promising platform for CNS-targeted TNFα inhibition and demonstrate a generalizable strategy for delivering biologics across the BBB with therapeutic-level exposure.

## Introduction

Tumor necrosis factor-alpha (TNFα) is a pleiotropic cytokine involved in neuroimmune signaling and synaptic modulation in the central nervous system (CNS). Under physiological conditions, TNFα is primarily produced by microglia, with additional contributions from astrocytes, where it plays important roles in glial signaling and the regulation of neuronal synaptic function. However, sustained elevation of TNFα promotes chronic neuroinflammation, disrupts synaptic plasticity, and has been implicated in the pathogenesis of several neurodegenerative disorders, including Alzheimer’s disease (AD). TNF-α levels are significantly elevated in the blood, cerebrospinal fluid (CSF), and CNS of a substantial subset of AD patients. This was first reported by Fillit et al. and later confirmed by multiple studies linking elevated proinflammatory cytokine levels to AD progression (1–6).

Beyond its immunomodulatory functions, TNFα critically influences synaptic physiology by promoting AMPA receptor (AMPAR) trafficking to the synaptic membrane and reducing synaptic GABA_A_ receptor (GABA_A_R) availability. This bidirectional effect disrupts the excitatory/inhibitory (E/I) balance, tipping it toward excitation, which increases neuronal activity and may exacerbate vulnerability to neurodegeneration (7–9). Aberrant excitatory transmission is known to promote the production of amyloid-beta (Aβ) and the secretion of tau protein— pathological hallmarks of AD (10–19). Thus, TNF-α, by modulating synaptic receptor dynamics and E/I balance, may act as an upstream driver of AD-related neuropathology.

Microglia, the principal source of TNF-α in the brain, are genetically linked to late-onset AD (LOAD) through mutations in *TREM2*, a microglia-specific receptor involved in immune signaling and phagocytosis (2, 20–22). Pathogenic variants such as TREM2 p.R47H impair microglial functions (23). Work from our group has shown that *Trem2*R47H knock-in (KI) rats recapitulate early LOAD phenotypes, including elevated CNS and CSF TNFα, increased excitatory transmission, reduced synaptic inhibition, and impaired long-term potentiation (LTP)—a cellular correlate of learning and memory (24–26). Notably, neutralizing TNFα in these models restores LTP and corrects synaptic dysfunction, directly implicating TNF-α as a key mediator of early AD-related pathophysiology.

Supporting this role, polymorphisms in *TNF* and *TNFR* genes have been associated with altered AD risk (27, 28) and experimental TNF-α blockade has been shown to reduce Aβ and tau pathology in preclinical models (29, 30). Epidemiological studies also report that chronic TNFα inhibitor (TNFI) use in patients with autoimmune conditions is associated with a reduced incidence of AD (31–33). Although anecdotal and underpowered, early clinical case studies have suggested symptomatic benefits from intrathecal or peri spinal TNFα inhibition in AD patients (34–36).

Despite their potent peripheral efficacy, FDA-approved TNFα inhibitors such as infliximab and etanercept exhibit negligible blood-brain barrier (BBB) permeability, with reported CSF-to-serum ratios around 0.001, severely limiting their utility for CNS disorders (6, 37–40). To overcome this limitation, we developed the NN-800 series of humanized nanobody-based biologics. NN-800s are heterotrimeric constructs comprising two high-affinity TNFα-neutralizing nanobodies (TNFI-α or TNFI-β) (41) and one humanized nanobody targeting the human transferrin receptor 1 (TfR1b-A2, -D1, -D2, or -D3) (42). By engaging TfR1, NN-800s leverage receptor-mediated transcytosis to cross the BBB—a strategy increasingly explored in CNS drug development (43–48).

Here, we demonstrate that NN-800s are the first TNFα inhibitors to combine high neutralizing potency with efficient CNS delivery. These dual properties position NN-800s as promising candidates for CNS-targeted TNFα inhibition in Alzheimer’s disease and other disorders characterized by supraphysiological TNFα levels.

## Results

### Heterotrimeric design preserves TfR1 binding and enhances TNFα inhibitory activity

To enable simultaneous BBB transport and TNFα neutralization, we combined two classes of humanized camelid nanobodies: (i) anti– TfR1 nanobodies capable of mediating BBB transcytosis (NewroBus; TfR1b-A2, -D1, -D2, and -D3), which correspond to NanoNewron candidates NN-104, NN-102, NN-103, and NN-101, respectively (42) and (ii) high-affinity TNFα-neutralizing nanobodies TNFI-α and TNFI-β, designated by NanoNewron as NN-223 and NN-224, respectively (IC_50_ = 48.84 pM and 64.31 pM) (41). As an initial step, eight heterodimeric constructs were generated by fusing each NewroBus variant to either TNFI-α or TNFI-β. These heterodimers retained both TfR1 binding and TNFα inhibitory activity, validating the NewroBus module as a functional BBB shuttle in bifunctional formats (42).

Because soluble TNFα exists as a homotrimer, we hypothesized that incorporation of two TNFα-binding domains within a single molecule would increase inhibitory potency through avidity-driven engagement of the trimeric ligand. To test this, we generated **Table 1**. TNFα inhibitory potency (IC_50_) of monomeric, heterodimeric, and heterotrimeric nanobodies. Intercalated heterotrimers show heterotrimeric constructs containing two TNFI nanobodies and one markedly enhanced potency. 96 TfR1-binding nanobody, initially using TfR1b-D3 (NN-101) as the BBB shuttle component (Figure 1A). All heterotrimers preserved TfR1 binding (Figure 1B), indicating that increasing valency did not impair interaction with the BBB transport receptor. Functionally, heterotrimers exhibited substantially enhanced TNFα neutralization compared with the corresponding heterodimers (Figure 1C). For example, the IC_50_ values of TNFI-α~TNFI-α~TfR1b-D3 and TNFI-β~TNFI-β~TfR1b-D3 were reduced by 4.3- and 4.9-fold, respectively, relative to TNFI-α~TfR1b-D3 and TNFI-β~TfR1b-D3. Notably, nanobody arrangement within the trimer had a major impact on activity. Intercalated configurations, in which the TfR1-binding domain was positioned between the two TNFI domains (TNFI-α~TfR1b-D3~TNFI-α and TNFI-β~TfR1b-D3~TNFI-β), showed the greatest potency, achieving 21-fold and 16.7-fold improvements, respectively, over the corresponding heterodimers.

**Table 1.**
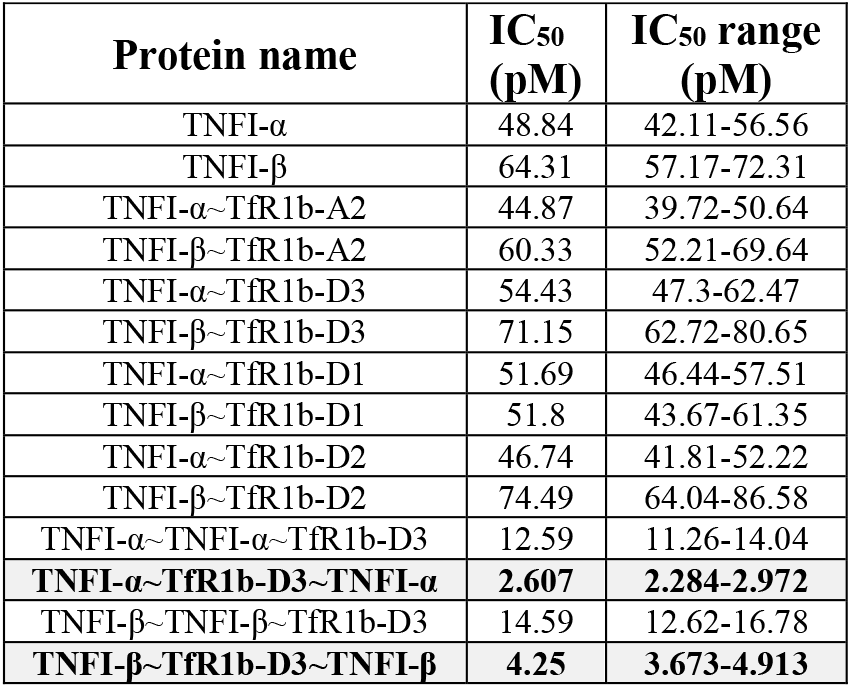
TNFα inhibitory potency (IC_50_) of monomeric, heterodimeric, and heterotrimeric nanobodies. Intercalated heterotrimers show markedly enhanced potency.

**Figure 1.**
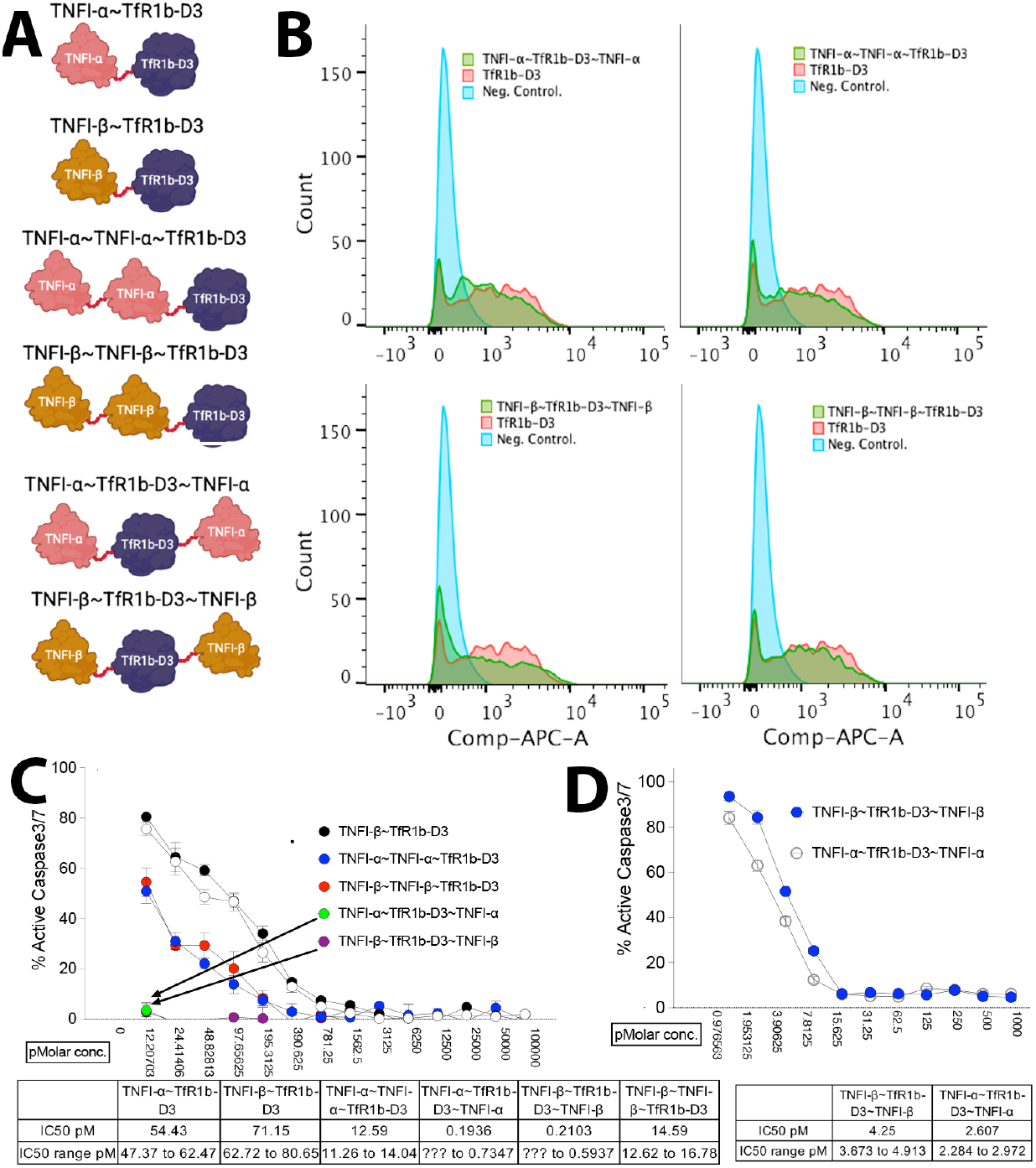
Heterotrimeric nanobodies enhance TNFα inhibition while preserving TfR1 binding. (A) Schematic of TfR1b-D3, TNFI8-α5, and TNFI-β constructs and corresponding heterodimeric and heterotrimeric formats. (B) Binding of TfR1b-D3 and TfR1b-D3–bas8e6d heterotrimers to HEK293 cells expressing human TfR1. (C) Dose-response inhibition of TNFα-induced apoptosis in WEHI-13VAR cells by heterodimers and heterotrimers. (D) IC_50_ values of heterotrimers from independent assays. Data are mean ± SEM (n=3).

Together, these results show that heterotrimeric architecture enhances TNFα neutralization while preserving TfR1 engagement and identify the intercalated format as the optimal configuration for combining BBB transport with high inhibitory activity (Table 1).

### Efficient in vivo BBB transcytosis of heterotrimeric nanobodies via TfR1 targeting

To evaluate in vivo BBB permeability, we selected TNFI-β~TfR1b-D3~TNFI-β and tested it in humanized Tfr1 knock-in (KI) rats, in which the endogenous rat *Tfrc* gene is replaced by the human *TFRC* sequence (49). This model enables expression of human TfR1, which is required for NewroBus binding, as these nanobodies do not recognize the rodent receptor (49). Accordingly, this system provides a physiologically relevant and necessary platform to assess TfR1-mediated CNS delivery in vivo.

Rats (Table 2) were intravenously injected with TNFI-β~TfR1b-D3~TNFI-β (1 µL/g of a 40 µM solution in PBS). Forty-eight hours post-injection, animals were perfused to remove circulating blood, and tissues were collected. Brains were further fractionated into vascular and parenchymal compartments using a protocol adapted from antibody transport vehicle (ATV) studies (50).

**Table 2.** genotype,sex and age of rats used in Figure 2 experiments.

| *TfR1* | sex | Age in days |
| --- | --- | --- |
| w/w | f | 99 |
| w/w | f | 112 |
| w/w | m | 100 |
| w/w | m | 99 |
| h/h | m | 112 |
| h/h | m | 112 |
| h/h | f | 112 |
| h/h | f | 112 |

TNFI-β~TfR1b-D3~TNFI-β was readily detected in whole brain homogenates, vascular fractions, and parenchymal fractions, indicating successful BBB transcytosis and tissue penetration (Figure 2a). CNS exposure was strictly dependent on human TfR1 expression, as no signal was detected in wild-type (*Tfr1*^*w/w*^) animals.

**Figure 2.**
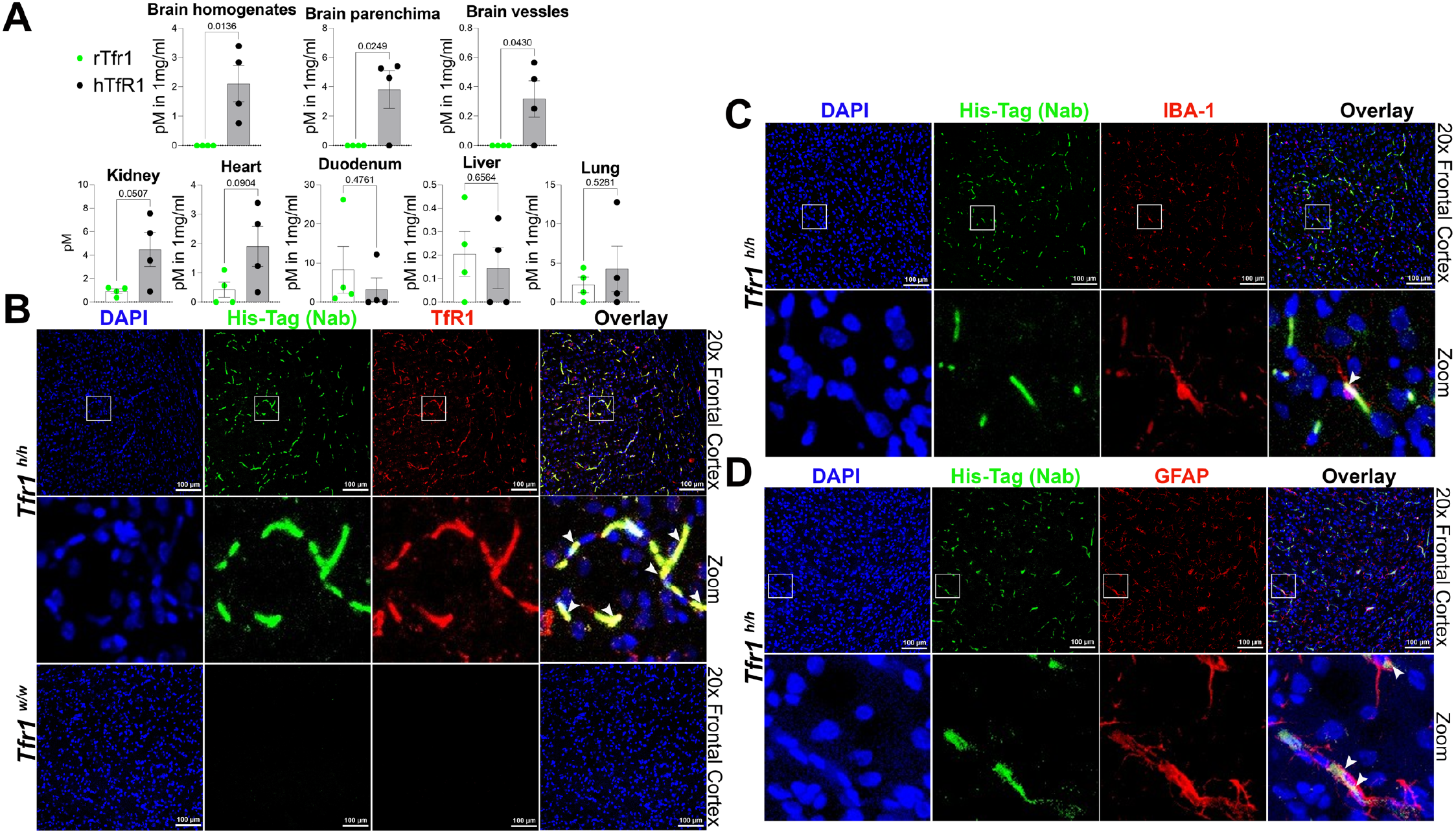
TNFI-β~TfR1b-D3~TNFI-β crosses the BBB via human TfR1. (A) Quantification of TNFI-β~TfR1b-D3~TNFI-β acros tissues. CNS exposure (brain homogenate, parenchymal and vascular fractions) was dependent on hTfR1 expression and observed only in *Tfr1*^*h/h*^ rats. CSF/serum ratios ranged from 0.123 to 0.26 (mean 0.17). (B) Immunofluorescence showing TNFI-β~TfR1b D3~TNFI-β (green, anti-His), hTfR1 (red), and DAPI in *Tfr1*^*h/h*^ but not *Tfr1*^*w/w*^ brains; arrows indicate colocalization. (C, D Colocalization of TNFI-β~TfR1b-D3~TNFI-β (green, anti-His) with IBA1-positive microglia (C) and GFAP-positive astrocytes (D) White arrows in the zoomed-in panel indicate the colocalization yellow spots within the region highlighted by the white square in th overlay panel.

Peripheral tissue distribution was limited, with only modest human TfR1–dependent accumulation observed in heart and kidney (Figure 2a). In contrast, brain uptake was robust, consistent with the high expression of TfR1 in brain endothelial cells and supporting preferential CNS targeting of NewroBus-based constructs. In addition, serum levels of the heterotrimer were consistently higher in humanized animals relative to wild-type controls, suggesting target-mediated stabilization, potentially via TfR1 binding and recycling.

Immunofluorescence analysis further confirmed CNS delivery and cellular distribution. TNFI-β~TfR1b-D3~TNFI-β was detected in the brains of *Tfr1*^*h/h*^ rats but not *Tfr1*^*w/w*^ controls (Figure 2b–d). The nanobody colocalized with hTfR1, indicating localization at brain vasculature. Notably, signal was also observed in IBA1-positive microglia and GFAP-positive astrocytes, demonstrating penetration beyond the vascular compartment and uptake into parenchymal cells.

### TNFI-β~TfR1b-D3~TNFI-β does not induce hematological toxicity in humanized Tf/TfR1 rats

Disruption of transferrin (TF)–TfR1 interactions or cellular iron uptake can result in hematological toxicity, particularly anemia. Consistent with this, homozygous TfR1 knockout mice are embryonically lethal, and hypo-transferrinemic mice develop severe anemia (51). Although in vitro studies showed that TfR1b-A and TfR1b-D nanobodies do not interfere with TF binding or uptake (42), we assessed the hematological safety of TNFI-β~TfR1b-D3~TNFI-β *in vivo*.

Complete blood count (CBC) analysis was performed in ~4-month-old rats humanized for both TF and TfR1. Animals received four subcutaneous doses of TNFI-β~TfR1b-D3~TNFI-β or vehicle (PBS) according to the schedule in Table 3. Baseline CBC values were obtained at Day −3, followed by measurements at 24 hours (after one dose), Day 17 (after three doses), and Day 24 (after four doses), enabling evaluation of both acute and cumulative effects.

Across all time points, hematological parameters remained within normal physiological ranges, with no significant differences compared to PBS-treated controls (Figure 3). All components of the white blood cell (WBC) profile—including total WBC counts, differential counts (neutrophils, lymphocytes, monocytes, eosinophils, and basophils), and their relative distributions—were unchanged. Similarly, red blood cell (RBC) parameters, including RBC count, hemoglobin (HGB), hematocrit (HCT), and erythrocyte indices (MCV, MCH, MCHC, RDW), showed no alterations, indicating absence of anemia or impaired erythropoiesis. Platelet parameters, including platelet count (PLT) and mean platelet volume (MPV), were also unaffected.

**Figure 3.**
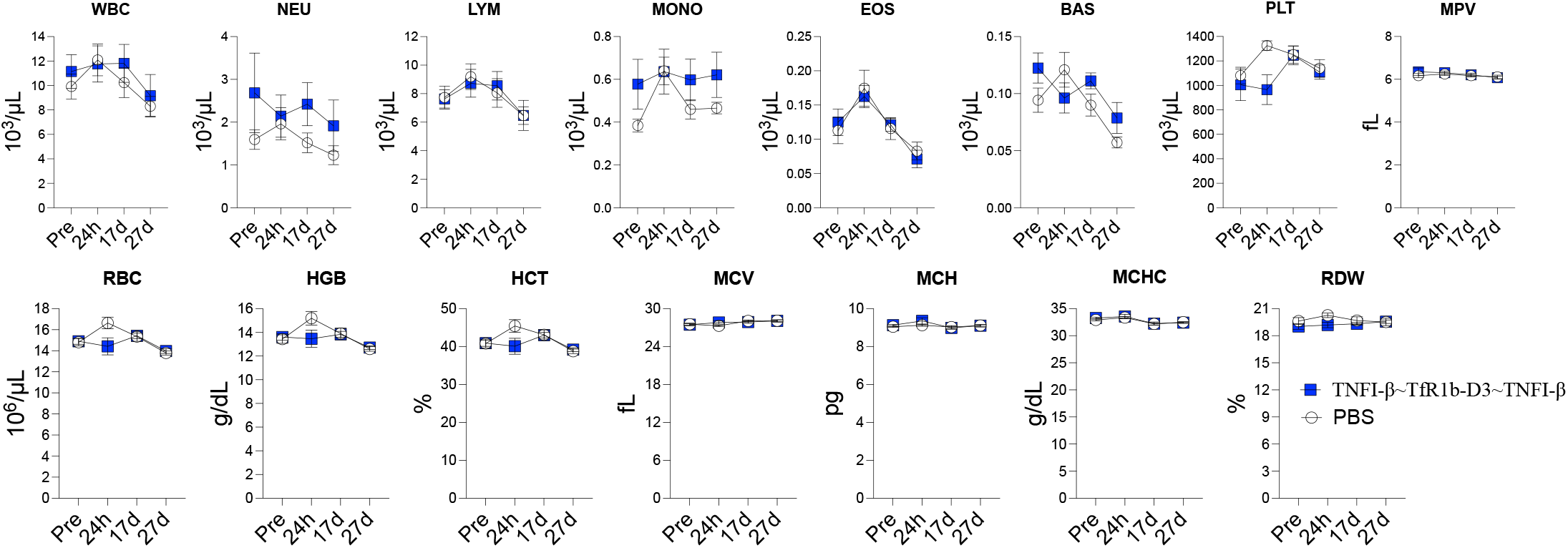
TNFI-β~TfR1b-D3~TNFI-β does not induce anemia in vivo. CBC analysis in ~4-month-old rats humanized for transferrin and TfR1 (*Tf*^*h/w*^*:Tfr1*^*h/h*^) following subcutaneous administration of TNFI-β~TfR1b-D3~TNFI-β or PBS. Measurements were performed at baseline (Day −3), Day 1, Day 17, and Day 24. No significant changes were observed across white blood cell, platelet or red blood cell parameters. These data indicate no detectable hematological toxicity or disruption of TF–TfR1–mediated iron homeostasis.

These findings indicate that TNFI-β~TfR1b-D3~TNFI-β does not impair TF–TfR1–mediated iron homeostasis or induce detectable hematological toxicity in vivo, even after repeated dosing. Given that NewroBus nanobodies selectively bind human TfR1 and that this model recapitulates human TF–TfR1 interactions, these data are expected to be predictive of clinical safety with respect to iron metabolism. Notably, the doses tested here exceed those anticipated for therapeutic use, further supporting a favorable safety margin.

Together, these results demonstrate a favorable tolerability profile for TNFI-β~TfR1b-D3~TNFI-β in a physiologically relevant humanized model.

### Efficient production and purification of heterotrimeric nanobodies

Based on the above findings, we generated eight intercalated heterotrimers by combining either TNFI-α or TNFI-β with each of the four NewroBus nanobodies (Figure 4A). All constructs were expressed in suspension-adapted CHO-S cells to ensure proper folding, disulfide bond formation, and compatibility with established biologics manufacturing platforms.

**Figure 4.**
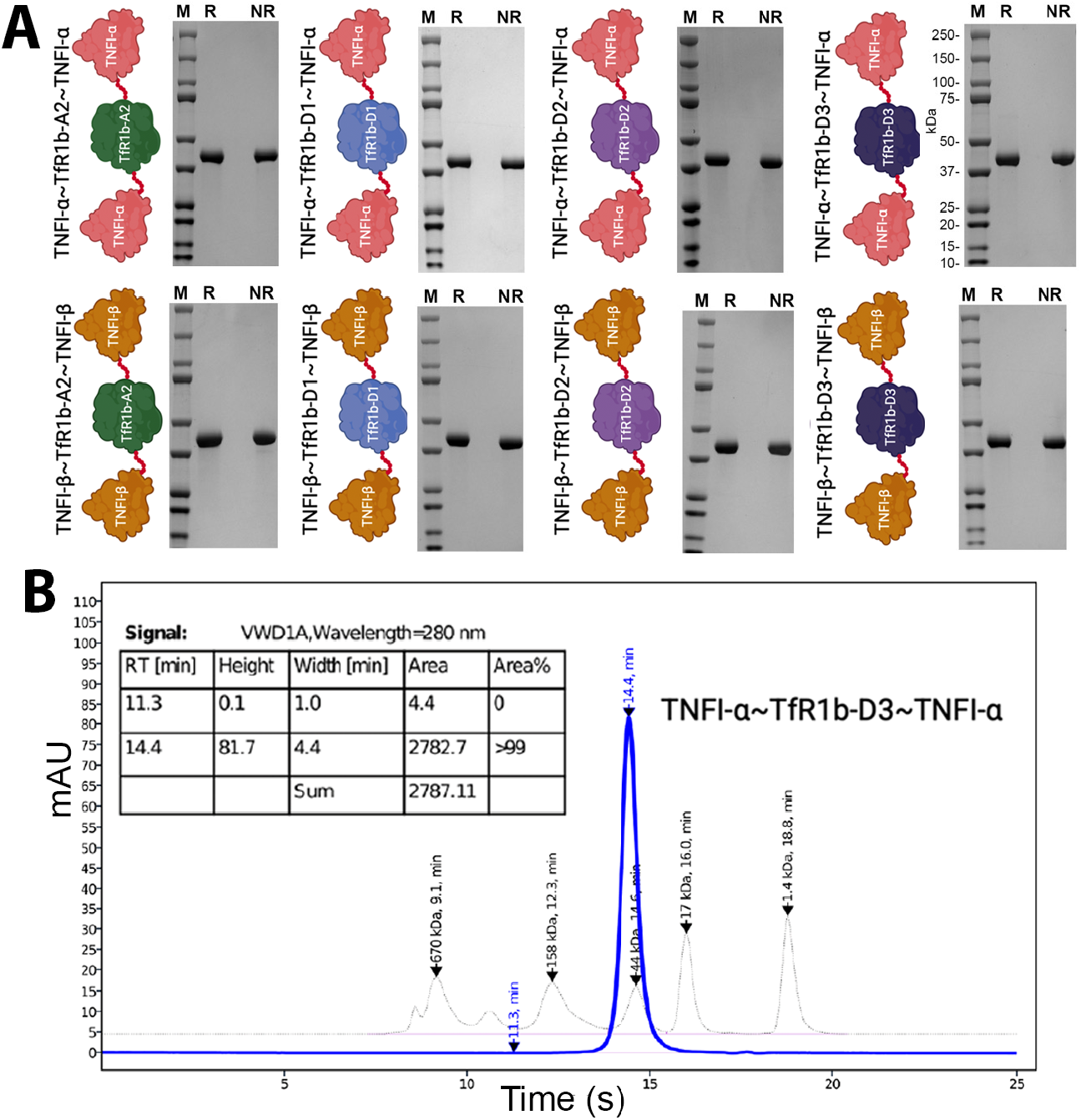
Production and purification of heterotrimeric nanobodies. (A) Schematic of the eight intercalated heterotrimers combining TNFI-α or TNFI-β with NewroBus variants and SDS-PAGE analysis under reducing (R) and non-reducing (NR) conditions. (B) Representative SEC-HPLC profile demonstrating high purity and minimal aggregation.

Transient transfections were performed at the 100 mL scale, and expressed proteins were purified using a two-step protocol consisting of immobilized metal affinity chromatography (IMAC) via a C-terminal His tag, followed by size-exclusion chromatography (SEC) for removal of aggregates and high-molecular-weight species. This approach yielded highly homogeneous products across all constructs.

All eight heterotrimers exhibited favorable biochemical and developability properties (Figure 4A,B; Table 4). Purity exceeded 99% by SEC-HPLC and 95% by SDS-PAGE, and endotoxin levels were consistently below 0.3 EU/mg. Final concentrations ranged from 0.69 to 1.6 mg/mL (17–40 µM), with total yields up to 67 mg per 100 mL culture.

**Table 4.** Expression yield and quality attributes of heterotrimeric nanobodies produced in CHO-S cells. All constructs were purified by IMAC followed by SEC. Proteins exhibited high purity (>99% SEC-HPLC), low endotoxin levels (≤0.3 EU/mg), and yields up to 67 mg per 100 mL culture.

| Construct (Long name) | NN ID | Conc. (mg/mL) | Purity SDS-PAGE (%) | Purity SEC-HPLC (%) | Endotoxin (EU/mg) | Yield (mg) | pI | MW (Da) | Conc. (μM) |
| --- | --- | --- | --- | --- | --- | --- | --- | --- | --- |
| TNFI-α~TfR1b-A2~TNFI-α | NN-844 | 1.10 | 95 | >99 | 0.045 | 53.9 | 8.24 | 39,526 | 27.8 |
| TNFI-α~TfR1b-D1~TNFI-α | NN-846 | 1.08 | 95 | >99 | 0.046 | 42.9 | 8.68 | 39,843 | 27.1 |
| TNFI-α~TfR1b-D2~TNFI-α | NN-848 | 0.69 | 95 | >99 | 0.236 | 15.5 | 8.68 | 39,839 | 17.3 |
| TNFI-α~TfR1b-D3~TNFI-α | NN-842 | 1.08 | 95 | >99 | 0.046 | 43.0 | 8.85 | 39,870 | 27.1 |
| TNFI-β~TfR1b-A2~TNFI-β | NN-843 | 1.40 | 95 | >99 | 0.036 | 58.8 | 8.24 | 39,528 | 35.4 |
| TNFI-β~TfR1b-D1~TNFI-β | NN-845 | 1.55 | 95 | >99 | 0.052 | 65.1 | 8.68 | 39,845 | 38.9 |
| TNFI-β~TfR1b-D2~TNFI-β | NN-847 | 1.60 | 95 | >99 | 0.301 | 67.2 | 8.68 | 39,841 | 40.2 |
| TNFI-β~TfR1b-D3~TNFI-β | NN-841 | 1.31 | 95 | >99 | 0.038 | 65.5 | 8.85 | 39,872 | 32.9 |

**Table 5.** Animal characteristics for subcutaneous PK/BBB studies. Human TfR1 knock-in (*Tfr1*^*h/h*^) rats (n=4per construct; 2 females, 2 males) were analyzed 24 h post-injection.

| NN ID | n<br>(F/M) | Age<br>(days) | Time<br>(h) |
| --- | --- | --- | --- |
| NN-844 | 4 (2/2) | 30 | 24 |
| NN-843 | 4 (2/2) | 30 | 24 |
| NN-846 | 4 (2/2) | 31 | 24 |
| NN-845 | 4 (2/2) | 31 | 24 |
| NN-848 | 4 (2/2) | 34 | 24 |
| NN-847 | 4 (2/2) | 34 | 24 |
| NN-842 | 4 (2/2) | 37–57 | 24 |
| NN-841 | 4 (2/2) | 37–68 | 24 |

These results demonstrate that heterotrimeric nanobodies can be efficiently produced using standard mammalian expression systems, with yields and quality attributes consistent with therapeutic development. Given the high expression levels observed under transient conditions, further scale-up using stable CHO cell lines is expected to substantially increase production yields and support scalable manufacturing.

Overall, the NN-800 heterotrimers display strong manufacturability characteristics—including high yield, high purity, low endotoxin content, and compatibility with established CHO-based production workflows—supporting advancement toward IND-enabling studies.

### Heterotrimers exhibit potent TNFα inhibition and cross-species activity with implications for preclinical models

All eight intercalated heterotrimers demonstrated potent TNFα-neutralizing activity, with IC_50_ values in the low single-digit picomolar range (Figure 5). Independent protein preparations yielded comparable results, confirming reproducibility. For example, NN-842 exhibited IC_50_ values of 2.61 pM (2.28–2.97) and 4.09 pM (3.13– 5.33) across batches, while NN-841 showed IC_50_ values of 4.25 pM (3.67–4.91) and 4.59 pM (3.57–5.90). These data demonstrate consistent TNFα inhibition across independently produced heterotrimers.

**Figure 5.**
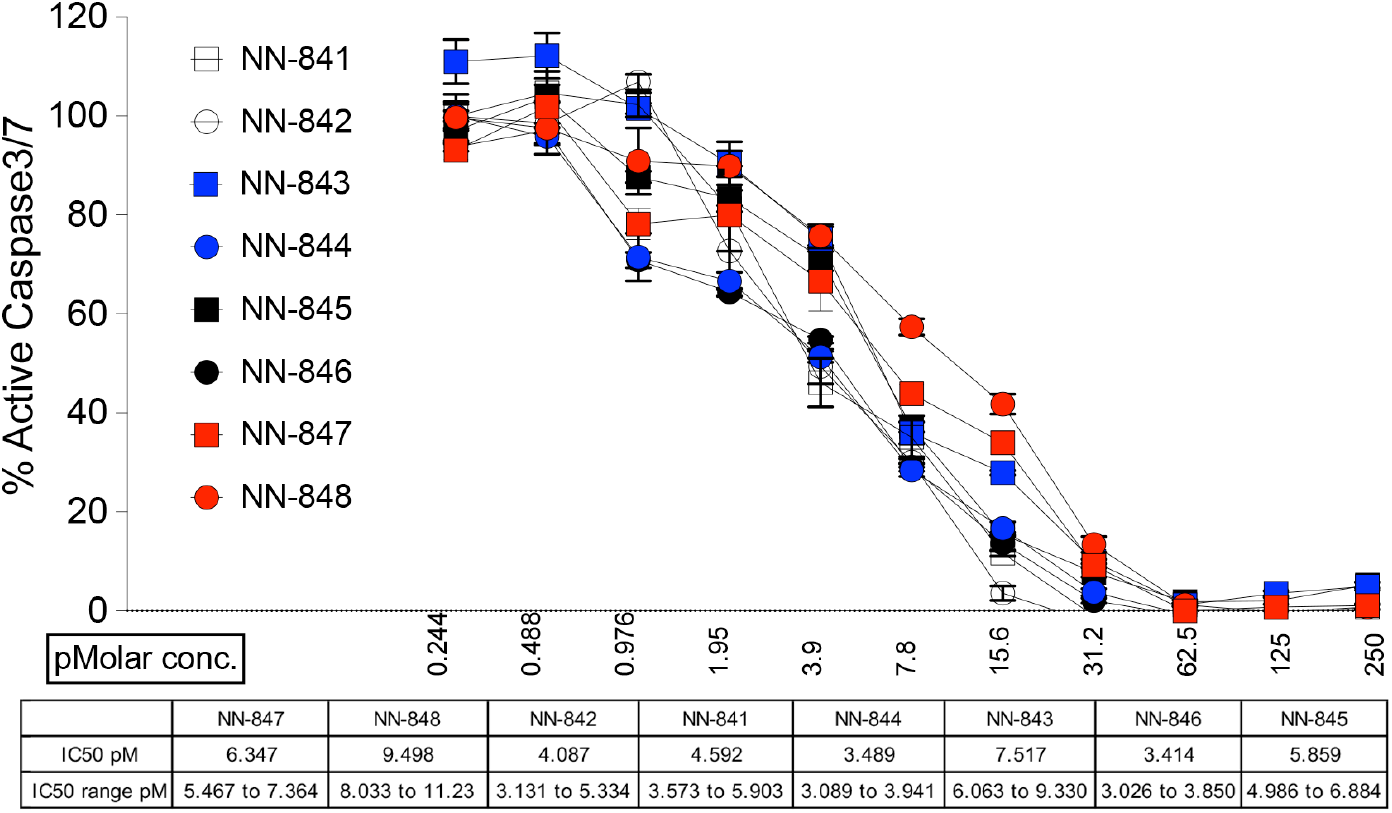
NN-800 heterotrimers exhibit potent TNFα-neutralizing activity. Dose-response inhibition of TNFα-induced apoptosis in WEHI-13VAR ce1l8ls7by eight heterotrimers (NN-841–NN-848). All constructs displayed IC_50_ values in the single-digit picomolar range. Data are mean ± SEM (n=4).

To assess cross-species activity, the TNFα-neutralizing potency of NN-800 heterotrimers (NN-841–NN-848) was evaluated against recombinant TNFα from rhesus macaque, rat, and mouse (Figure 6). All constructs retained potent neutralizing activity against rhesus macaque TNFα, with IC_50_ values in the single-digit picomolar range, comparable to those observed for human TNFα, indicating conservation of the targeted epitope across primates.

**Figure 6.**
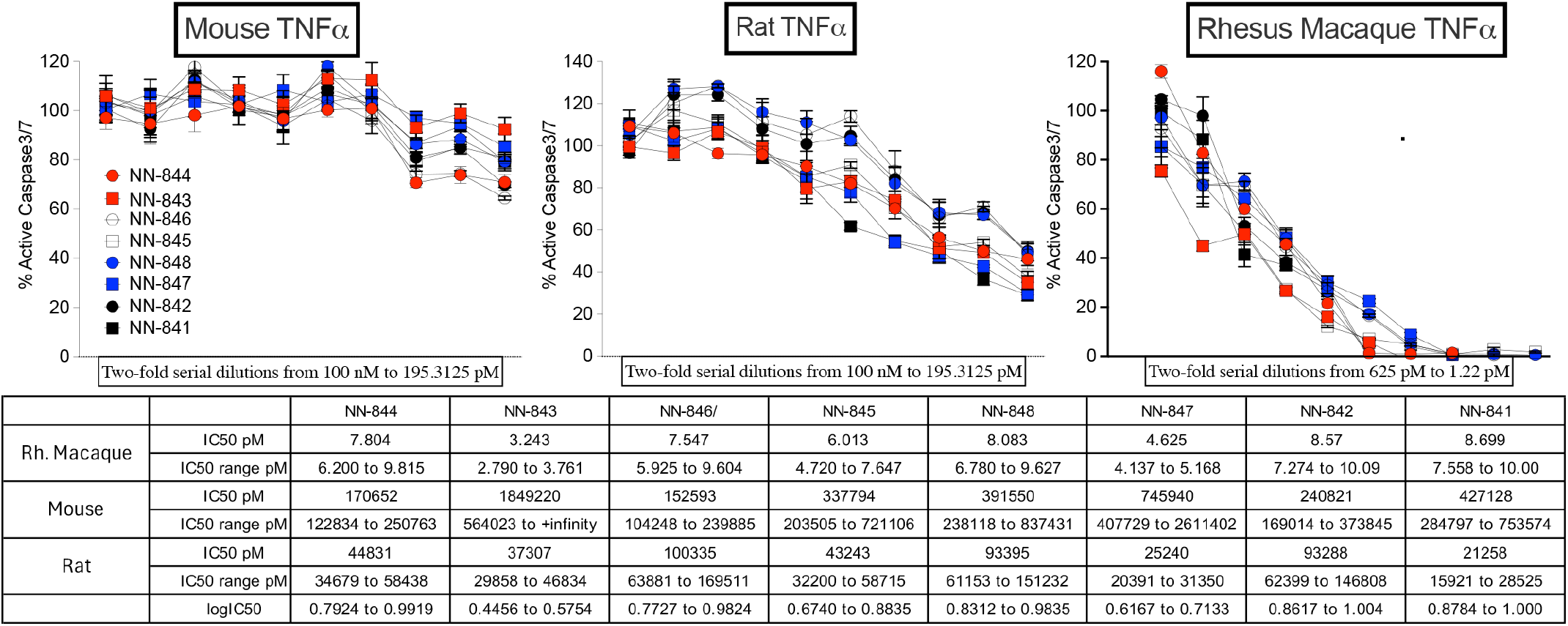
Cross-species TNFα inhibition by NN-800 heterotrimers. Dose-response inhibition of TNFα from rhesus macaque, rat, and mouse by NN-800 heterotrimers (NN-841–NN-848). All constructs exhibited potent activity against rhesus TNFα (single-digit pM IC_50_), reduced potency against rat TNFα (~10,000-fold), and minimal or no activity against mouse TNFα. Data are mean ± SEM (n=4).

In contrast, activity against rodent TNFα was markedly reduced. IC_50_ values against rat TNFα shifted to the double-digit nanomolar range (~10,000-fold reduction in potency), and minimal or no activity was detected against mouse TNFα, consistent with prior characterization of TNFI-α and TNFI-β and reflecting sequence divergence across species.

Notably, these assays were performed using protein preparations stored at −80 °C for up to 6.5 months, including samples subjected to a freeze–thaw cycle. All constructs retained full activity against rhesus TNFα, indicating strong stability under storage conditions relevant for preclinical development.

These findings have important implications for model selection. The lack of activity against rodent TNFα precludes evaluation of TNFα inhibition in standard mouse or rat models unless TNFα is humanized. Conversely, although nonhuman primates are suitable for assessing TNFα engagement, the NewroBus modules do not cross-react with nonhuman primate TfR1, precluding evaluation of BBB transport in these species. Together, these constraints highlight the need for humanized models to assess both target engagement and CNS delivery.

### Heterotrimers exhibit efficient BBB penetration following subcutaneous administration

We next evaluated BBB penetration following subcutaneous (SQ) administration in. We next tested whether these heterotrimers can cross the BBB following subcutaneous (SQ) injection, a clinically relevant route for patient-administered biologics due to its convenience, safety, and cost-effectiveness.Compared to intravenous administration, SQ delivery enables at-home dosing (e.g., Humira, Enbrel, Simponi), provides gradual absorption and sustained systemic exposure, and reduces the need for clinic-based infusions, thereby improving patient compliance and overall treatment burden.

*Tfr1*^*h/h*^ rats (n=4 per construct; 2 males, 2 females) received 1 µL/g body weight of a 17.3 µM solution of each heterotrimer. Serum and cerebrospinal fluid (CSF) concentrations were measured 24 hours post-injection (Figure 7). All constructs achieved detectable systemic and CNS exposure. Serum concentrations ranged from 50.2 pM (NN-841) to 390 pM (NN-844), while CSF concentrations ranged from 18.3 pM (NN-841) to 60 pM (NN-844), confirming BBB penetration for all heterotrimers. CSF/serum ratios ranged from 0.14 (NN-843 and NN-846) to 0.40 (NN-842), consistent with efficient TfR1-mediated transcytosis. NN-842 exhibited the highest CNS delivery efficiency (CSF/serum = 0.40), whereas NN-844 reached the highest absolute concentrations in both serum and CSF.

**Figure 7.**
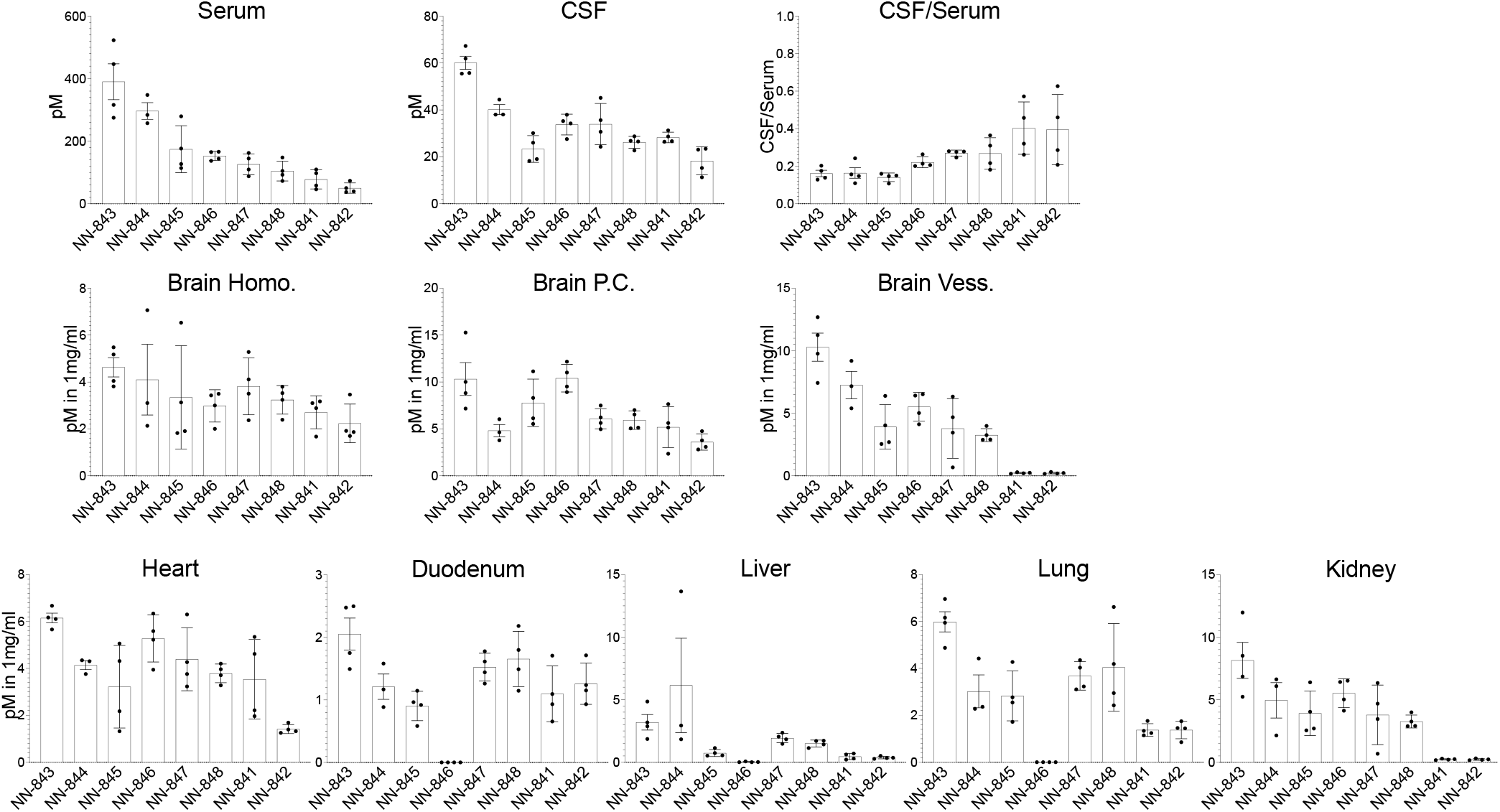
Tissue distribution of NN-800 heterotrimers 24 h after subcutaneous administration in *Tfr1*^*h/h*^ rats, including serum CSF, brain fractions (homogenate, parenchymal, vascular), and peripheral organs.

The observed variability in CNS exposure likely reflects differences in TfR1 binding, molecular architecture, and intracellular trafficking among constructs. Overall, the distribution profile following SQ administration was consistent with intravenous delivery (Figure 2) and prior heterodimer data (42).

To further assess cellular distribution within the CNS, we performed immunofluorescence analysis of brain sections following administration of NN-800 heterotrimers. Representative data are shown for NN-841 (Figure 8). Consistent with the results shown in Figure 2, NN-841 was detected in both vascular and parenchymal compartments in Tfr1^h/h^ rats, with clear colocalization with hTfR1 in brain vessels.

**Figure 8.**
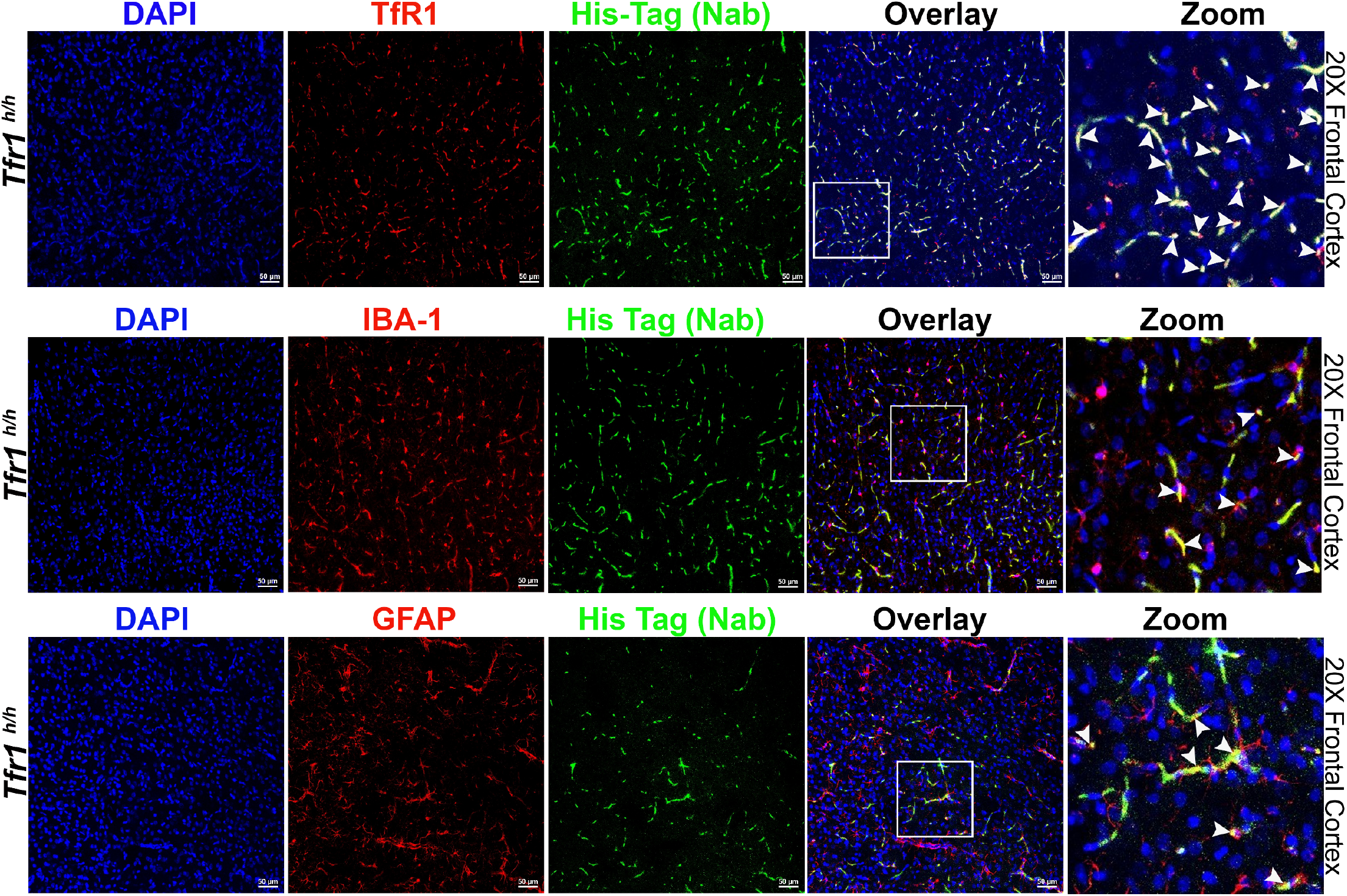
Cellular localization of NN-800 heterotrimers in the brain. Representative immunofluorescence of NN-841 in *Tfr1*^*h/h*^ rat brain. NN-841 (green, anti-His), hTfR1 (red), and DAPI show signal in brain vasculature and parenchyma; arrows indicate colocalization (top). NN-841 is detected in IBA1-positive microglia and GFAP-positive astrocytes (middle and bottom). White arrows in the zoomed-in panel indicate the colocalization yellow spots within the region highlighted by the white square in the overlay panel.

Importantly, NN-841 signal was also observed in IBA1-positive microglia and GFAP-positive astrocytes, indicating that the heterotrimer not only crosses the BBB but is internalized into resident CNS cells including cell types implicated in TNFα-driven neuroinflammation. This distribution beyond the vascular compartment supports effective delivery to biologically relevant target cells within the brain.

Similar localization patterns were observed for all NN-800 heterotrimers (Supplementary Figures 1–7), demonstrating that CNS penetration and cellular uptake are consistent features of this class of molecules.

Together, these results demonstrate that NN-800 heterotrimers retain high TNFα inhibitory potency and achieve robust CNS exposure following subcutaneous administration, supporting their suitability for clinically relevant dosing strategies.

## Discussion

Biologic TNFα inhibitors—such as adalimumab, etanercept, and infliximab—are potent and specific therapeutics with well-established efficacy in systemic inflammatory diseases. Observational studies have reported a reduced incidence of AD among patients receiving these therapies for peripheral conditions, despite their minimal BBB penetration (CSF/serum ~0.001) (31–33).

In contrast, XPro™1595 has shown inconsistent efficacy in Phase II trials for AD, despite promising results in preclinical models (29). This discrepancy may be explained by substantially lower dosing in human studies (approximately 50-fold lower than in animal models), limited CNS penetration (CSF/serum ~0.001, ~4000-fold lower than NN-800s) (52), and lower TNFα neutralizing potency (IC50 ~1–5 nM, ~1000-fold less potent than NN-800s). Collectively, these factors suggest that inadequate central TNFα inhibition may have contributed to the lack of clear clinical efficacy.

Increasing evidence supports a direct pathogenic role of TNFα in the CNS. One key mechanism is disruption of synaptic receptor trafficking and excitatory/inhibitory balance: TNFα enhances excitatory signaling by promoting GABA_A receptor internalization and AMPA receptor insertion, driving network hyperexcitability (7–9). This heightened activity is linked to increased production of amyloid-β and propagation of tau pathology, both of which are activity-dependent (10–19)

Elevated TNFα also induces maladaptive glial responses. Microglia shift to a pro-inflammatory, less phagocytic state, while astrocytes become reactive, with impaired glutamate uptake and compromised BBB integrity (53–63). Together, these changes sustain a cycle of neuroinflammation and neurodegeneration.

Human studies further support this role, including associations between TNF/TNFR polymorphisms and Alzheimer’s disease risk (27, 28), and small clinical studies suggesting symptomatic improvement with central TNFα inhibition (34–36).

These findings raise two key questions: (1) is TNFα a central driver of neuroinflammatory and neurodegenerative diseases, and therefore a viable target for disease-modifying therapies? (2) If so, do BBB-impermeable TNFα inhibitors act primarily through peripheral mechanisms, or could limited but sustained CNS exposure mediate their effects?

The NN-800 program was developed to address these questions. NN-800 molecules are engineered as heterotrimeric constructs composed of two high-affinity humanized TNFα-binding VHH nanobodies (41) flanking a central BBB carrier module, NewroBus (42). NewroBus was selected for its ability to bind transferrin receptor 1 (TfR1) without interfering with transferrin binding or iron uptake *(Fig. 3 and previous report* (42))—an important feature for maintaining physiological iron homeostasis. By engaging TfR1, NN-800 enables receptor-mediated transcytosis across the BBB while preserving a favorable safety profile.

In vivo data support this design: NN-800 trimers achieve high CNS exposure, with CSF-to-serum ratios approaching 0.5 following subcutaneous administration and accumulate in the brain at levels predicted to be therapeutically meaningful. Importantly, CNS drug delivery must be evaluated not only by CSF/serum ratios but also by absolute brain concentrations. While high ratios indicate efficient BBB transport, therapeutic efficacy ultimately depends on achieving sufficient CNS exposure. Compounds with more modest ratios may still be effective if systemic exposure is sufficient.

NN-800 molecules appear to balance these parameters by combining efficient BBB transport with sustained systemic exposure, enabling therapeutically relevant CNS concentrations while maintaining peripheral TNFα inhibition at levels that may reduce the risk of systemic immunosuppression. Modest peripheral TNFα modulation may also contribute to efficacy in diseases with both central and systemic inflammatory components.

Altogether, the NN-800 platform offers a promising solution for delivering TNFα inhibitors to the CNS. Its ability to combine high BBB permeability, strong TNFα neutralization, and a favorable safety profile supports its development for AD and other neuroinflammatory or neurodegenerative disorders characterized by supraphysiological TNFα levels.

Importantly, the data presented here establish robust CNS delivery, target engagement, and safety, but do not yet demonstrate disease-modifying efficacy in AD models. This will be a critical next step to confirm therapeutic potential.

Several aspects remain to be addressed to fully de-risk NN-800s for clinical development. Binding affinities for both TNFα and hTfR1 in the trimeric format have not yet been determined, and structural characterization will be important to distinguish contributions of affinity and avidity. In addition, while immunogenicity was minimized through design, ongoing ex vivo studies using human immune cells are required to confirm immune compatibility across HLA haplotypes.

In addition, rodent studies require humanized models to evaluate both TNFα inhibition and BBB transport, as NN-800s do not effectively inhibit rodent TNFα and NewroBus does not bind nonhuman primate TfR1. This species specificity is not incidental but reflects the selection strategy used to generate NewroBus nanobodies. Specifically, NewroBus variants were selected to bind human TfR1 without interfering with transferrin binding, receptor-mediated endocytosis, or iron transport. Notably, nanobodies that preserved these functions were consistently human-specific, whereas cross-reactive binders interfered with TfR1 physiological functions (42).

Because functional TfR1 domains involved in transferrin binding and trafficking are highly conserved across species, cross-reactivity often indicates binding to these regions, increasing the likelihood of disrupting TfR1 biology and causing toxicity. Thus, cross-reactivity, often considered advantageous for preclinical testing, may instead represent a potential safety liability in humans. In contrast, NewroBus nanobodies appear to recognize less conserved, non-functional regions of TfR1, enabling receptor engagement and transcytosis without perturbing iron homeostasis.

Importantly, assessment of BBB transport requires evaluation in the context of human TfR1, as receptor binding kinetics play a central role in transcytosis. The rates of association and dissociation (k_on and k_off), and thus the residence time of the ligand–receptor complex, critically influence intracellular trafficking, endosomal sorting, and release into the brain parenchyma. Even for TfR1-targeting shuttles that are cross-reactive across species, these kinetic parameters are unlikely to be identical between human and animal TfR1 due to subtle sequence and structural differences. As a result, a construct that exhibits efficient BBB transcytosis in rodents or nonhuman primates may display altered trafficking behavior in humans, including reduced transcytosis efficiency or increased lysosomal degradation.

Accordingly, evaluation of BBB penetration using human TfR1, such as our humanized Tfr1 rats, is essential for accurate prediction of clinical performance. Together with the safety considerations described above, these findings suggest that the lack of cross-species reactivity is not a limitation but rather a feature of functionally optimized TfR1 shuttles that preserve physiological TfR1 function.

In summary, the NN-800 platform represents a promising strategy for CNS-targeted TNFα inhibition, combining high potency, efficient BBB penetration, and a favorable safety profile.

## Materials and Methods

### Animals

All animal procedures were conducted in accordance with the NIH *Ethical Guidelines for the Treatment of Laboratory Animals*. All protocols were approved by the Rutgers Institutional Animal Care and Use Committee (IACUC Protocol #201,702,513). Efforts were made to minimize animal suffering and reduce the number of animals used.

### Cell lines

Cell lines used in this study are listed below:

| Cell Line | Description | Vendor | Catalog No. |
| --- | --- | --- | --- |
| WEHI-13VAR | Mouse fibrosarcoma TNFα-sensitive reporter cell line | ATCC | CRL-2168 |
| HEK-ATP089 | HEK293 cells stably expressing human Tfr1 | Acro Biosystems | CHEK-ATP089 |

### Primary and secondary antibodies

Antibodies used for flow cytometry, Western blotting, and immunohistochemistry are listed below:

| Target | Antibody | Vendor | Catalog No. |
| --- | --- | --- | --- |
| His tag | APC-conjugated | R&D Systems | IC050A |
| His tag | AF488 Rabbit mAb | Cell Signaling Technology | 14930 |
| VHH domain | Goat anti-alpaca | Jackson ImmunoResearch | 128-005-230 |
| Human Tfr1 | APC-conjugated | Invitrogen | 17071942 |
| Iba1 | Microglia marker | Wako | 012-28401 |
| GFAP | Astrocyte marker | BD Biosciences | 556327 |
| Goat IgG | SULFO-TAG | MSD | R32AG |
| Mouse IgG | Alexa Fluor 594 | Invitrogen | A-11032 |
| Rabbit IgG | HRP-conjugated | Cell Signaling Technology | 7074 |

### Recombinant proteins and reagents

Reagents used for binding, detection, and functional assays are summarized below:

**Recombinant proteins and reagents.** Reagents used for binding, detection, and functional assays are summarized below:
| Reagent | Description | Vendor | Catalog No. |
| --- | --- | --- | --- |
| Human TNFα | Recombinant | Acro Biosystems | TNA-H5228 |
| Biotinylated TNFα | ELISA detection | Acro Biosystems | TNA-H82E3 |

### General reagents and instrumentation

General reagents and instrumentation are listed below:

**General reagents and instrumentation.** General reagents and instrumentation are listed below:
| Reagent | Vendor | Catalog No. |
| --- | --- | --- |
| FBS | Gibco | A3840102 |
| DMEM | Corning | 10-017-CV |
| RPMI | Corning | 10-040-CV |
| Actinomycin D | Sigma | A9415 |
| Caspase-3/7 reagent | Thermo Fisher | C10423 |
| Dextran | Sigma | 31397 |
| Hygromycin B | Sigma | H3272 |

| Instrument | Vendor |
| --- | --- |
| MESO QuickPlex SQ 120 | MSD |
| Incucyte Live-Cell Analysis System | Sartorius |
| Leica Stellaris 8 tau STED | Leica |
| Leica CM1950 Cryostat | Leica |

### Protein expression and purification

Heterotrimeric nanobody constructs were expressed in suspension-adapted CHO-S cells using transient transfection at a 100 mL scale. All constructs contained a C-terminal 6×His tag to enable purification.

Cell supernatants were harvested and clarified prior to purification. Proteins were purified using immobilized metal affinity chromatography (IMAC) followed by size-exclusion chromatography (SEC) to remove aggregates and ensure homogeneity.

Purified proteins exhibited ≥95% purity by SDS-PAGE and >99% purity by SEC-HPLC, with endotoxin levels ≤0.3 EU/mg. Final concentrations ranged from 0.69 to 1.6 mg/mL (17–40 µM), with total yields up to 67 mg per 100 mL culture. Protein production was performed by GenScript.

### FACS analysis using stable HEK293-hTfR1 cells

HEK293-hTfR1 cells (Acro Biosystems, CHEK-ATP089), a stable clone expressing human TfR1, were maintained in DMEM (Corning, 10-017-CV) supplemented with 10% fetal bovine serum (Gibco, A3840102) at 37°C in a humidified 5% CO_2_ incubator. Cells were harvested for analysis 24–48 hours post-transfection.and used as a positive control for TfR1 surface expression and nanobody binding. 100μg/mL of Hygromycin B (Sigma, H3272) was used for specific selection of TfR1 expressed clones.

#### Staining procedure

Cells were resuspended in ice-cold FACS buffer (PBS containing 1% BSA and 1 mM EDTA, pH 7.2) and incubated with nanobodies (final concentration 40–100 nM) at 4°C for 45 minutes with gentle shaking. After three washes in FACS buffer, cells were incubated for 30 minutes at 4°C with APC-conjugated anti-His tag antibody (R&D Systems, IC050A) at 1:100 dilution to detect bound His-tagged nanobodies. Following additional washes, cells were stained with propidium iodide (Invitrogen, P3566; 1:1000 dilution) to exclude dead cells.

#### Flow cytometry analysis

Samples were analyzed on a BD LSR Fortessa or similar flow cytometer. Data were gated on live, EGFP-positive cells (for transfection marker) and analyzed for APC signal using FlowJo software.

### TNFα inhibition and IC_50_ determination using incucyte caspase-3/7 assays

WEHI-13VAR cells (ATCC, CRL-2168), a TNFα-sensitive mouse fibrosarcoma line, were cultured in RPMI 1640 medium (Corning, 10-040-CV) supplemented with 10% fetal bovine serum (Gibco, A3840102) at 37°C with 5% CO_2_. Cells were seeded into 96-well plates at 30,000 cells per well in RPMI containing 10% FBS and 1 μg/mL Actinomycin D (Sigma-Aldrich, A9415) to sensitize cells to TNFα-induced apoptosis. Recombinant human TNFα (Acro Biosystems, TNA-H5228) was pre-incubated with varying concentrations of TNFI nanobodies for 30 minutes at room temperature, then added to the cells at a final TNFα concentration of 0.25 ng/mL. Caspase-3/7 activity was monitored using Caspase-3/7 Green Detection Reagent (Thermo Fisher Scientific, C10423) added at a final concentration of 0.5 μM. Plates were immediately placed in the Incucyte Live-Cell Imaging System (Sartorius), and fluorescent signals were monitored every 2–4 hours for up to 24 hours. Fluorescent caspase-3/7 signal intensity was quantified using Incucyte analysis software. TNFα-induced apoptosis was set to 100%, and inhibition was expressed as a percentage of this maximum signal. IC_50_ values for each TNFI-Nb were calculated using GraphPad Prism with nonlinear regression (log[inhibitor] vs. response – variable slope).

### In vivo drug administration, perfusion, and sample collection. *Intravenous (IV) injection*

Rats were briefly warmed under a heat lamp to promote vasodilation. Intravenous drug administration was performed via the lateral tail vein using 1 mL Tuberculin Syringes fitted with 25G needles (Cardinal Health). Nanobodies or fusion constructs were injected slowly at a volume of 1 µL per gram of body weight in sterile PBS to minimize stress and ensure accurate dosing.

#### SQ injection

For SQ administration, animals were gently restrained using a towel, and the compound was injected into the interscapular area using a 1mL Tuberculin syringes fitted with 25G needles (Cardinal Health). Dosing volume was the same as IV: 1 µL of a 40 µM solution per gram of body weight in PBS. Animals were monitored for signs of discomfort or inflammation at the injection site.

#### Perfusion and tissues collection

At the designated time points post-injection, animals were deeply anesthetized using isoflurane. For terminal experiments involving brain collection, transcardiac perfusion was performed with cold PBS (without calcium or magnesium) at a flow rate of 10 mL/min for 10 minutes to remove blood from the vasculature. Heart, liver, lung, kidney, duodenum and brain were rapidly dissected post PBS perfusion and homogenized with a glass-Teflon homogenizer in 250 mM sucrose, 20 mM Tris-base pH 7.4, 1 mM EDTA, 1 mM EGTA plus protease, and phosphatase inhibitors.

### Serum collection

#### Non-terminal

Blood samples (~300 µL) were collected via tail vein using a 25G VACUETTE Safety Blood Collection Set (Greiner Bio-One) into BD Microtainer Serum Separator Tubes (Becton Dickinson). Samples were allowed to clot for 30 minutes at room temperature, followed by centrifugation at 9000 × g for 30 seconds.

#### Terminal

Blood was collected by cardiac puncture using a 5 mL syringe (Air-Tite Products Co., Inc) fitted with a 21G needle (Cardinal Health), transferred into BD Vacutainer SST tubes, allowed to clot for 30 minutes at room temperature, and centrifuged at 2000 × g for 10 minutes. All serum samples were aliquoted and stored at –80°C.

#### CSF collection

Following transcardiac perfusion and prior to brain tissue collection, CSF was collected via cisterna magna puncture. Animals were placed in a stereotaxic frame (Stoelting Co.). The head was flexed forward, and a midline incision was made at the base of the skull to expose the translucent dura mater over the cisterna magna, using a 28G insulin syringe (EXEL), CSF was slowly withdrawn, carefully avoiding blood contamination. Typically, 50–80 µL of CSF was collected per animal. Samples were snap-frozen in liquid nitrogen and stored at –80°C.

### ELISA for TNFα-based detection of nanobodies

#### TNFα-based ELISA

Streptavidin-coated 96-well plates (Meso Scale Discovery, L15SA or L45SA) were blocked overnight at 4°C with 3% BSA in PBST (PBS + 0.05% Tween-20). Plates were then coated with 0.25 µg/mL biotinylated human TNFα (Acro Biosystems, TNA-H82E3-25ug) in PBS for 1 hour at room temperature with shaking. After washing 4× with PBST, samples containing nanobodies were added and incubated overnight at 4°C. Following three PBST washes, wells were incubated with 1 µg/mL Goat Anti-Alpaca IgG, VHH domain (Jackson ImmunoResearch, 128-005-230) for 1 hour at room temperature. After additional washes, SULFO-TAG-labeled anti-goat IgG secondary antibody (Meso Scale Discovery, R32AG; 0.5–1 µg/mL) was added and incubated for 1 hour.

#### Plate reading and analysis

After final washes, wells were developed using 2× MSD Read Buffer (Meso Scale Discovery, R92TC) and read on a MESO QuickPlex SQ 120 instrument. Background-subtracted signals were normalized to control wells. TNFI-Nb1 (specific for TNFα) and irrelevant control nanobodies were used to confirm specificity.

### Hematotoxicity Assessment via Complete Blood Count (CBC)

To evaluate hematotoxicity following intravenous administration of TfR1-targeting nanobody constructs, complete blood counts (CBCs) were performed in treated and control rats at multiple time points. Animals received IV injections of either PBS (Group 1, n=7; 3 males, 4 females) or TNFI-β–TfR1b–A2 (Group 2, n=8; 3 males, 5 females) at a dose of 1 µL of a 40 µM solution per gram of body weight. Prior the blood collection, rats were briefly warmed under a heat lamp. Blood samples were collected from the tail lateral vein using a VACUTTE Safety Blood Collection Set (Greiner Bio-One) and transferred into MiniCollect K2EDTA tubes (Greiner Bio-One). Samples were immediately placed on an icepack and kept at +4°C until analysis and transported to the In Vivo Research Services (IRVS) Core Facility at Rutgers University. CBCs were performed using the Heska Element HT5 CBC Analyzer.

CBCs were collected at four time points:

- **Day –3 (D–3):** Baseline, prior to treatment
- **Day 1 (D1):** 24 hours after the first injection
- **Day 17 (D17):** After the third injection
- **Day 24 (D24):** After the fourth injection The following parameters were measured:

#### White blood cell (WBC) profile

- Total WBC (×10^3^/μL)
- Neutrophils (NEU), Lymphocytes (LYM), Monocytes (MONO), Eosinophils (EOS), Basophils (BAS)
- Percent distribution: NEU%, LYM%, MONO%, EOS%, BAS%

#### Red blood cell (RBC) profile

- RBC count (×10^6^/μL), Hemoglobin (HGB, g/dL), Hematocrit (HCT, %)
- Mean corpuscular volume (MCV, fL), Mean corpuscular hemoglobin (MCH, pg)
- Mean corpuscular hemoglobin concentration (MCHC, g/dL)
- Red cell distribution width (RDW, %)

#### Platelet profile

- Platelet count (PLT, ×10^3^/μL), Mean platelet volume (MPV, fL)

### Fractionation of brain tissue

To evaluate the distribution of nanobodies between the brain vasculature and parenchyma, rats were deeply anesthetized and perfused transcardially with cold PBS. The right hemisphere was dissected, and choroid plexuses were removed. Tissue was homogenized in 5 mL of ice-cold S1 buffer (250 mM sucrose, 20 mM Tris-base pH 7.4, 1 mM EDTA, 1 mM EGTA) using a glass-Teflon homogenizer (10 strokes). Homogenates were centrifuged at 1,000 × g for 10 minutes at 4°C. Pellets were resuspended in 2 mL of 17% dextran (MW 60,000; Sigma, 31397) and centrifuged at 4,200 × g for 15 minutes. The pellet was collected as the capillary-enriched vasculature fraction. The supernatant was diluted in S1 buffer and centrifuged again at 4,200 × g for 15 minutes. The resulting pellet was collected as the vascular-depleted parenchymal fraction.

Both fractions were lysed in ice-cold S1 buffer containing protease and phosphatase inhibitors and sonicated (50% amplitude, 3 × 10 s bursts with 30 s interval rests). Protein concentrations were determined by Bradford assay. Aliquots were used for ELISA and Western blot.

### Immunohistochemistry (IHC)

Rats were deeply anesthetized and perfused transcardially with PBS (without calcium and magnesium), followed by fixation with 4% paraformaldehyde (PFA; Electron Microscopy Sciences, 15714-S) in PBS at a flow rate of 10 mL/min for 10 minutes. Brains were extracted and post-fixed in 4% PFA at 4°C for 24 hours on a shaker. After fixation, brains were washed twice in PBS and incubated in 30% sucrose in PBS at 4°C for 48 hours for cryoprotection. Brains were embedded in OCT compound (Fisher, 23-730-571), frozen, and stored at –80°C. Coronal sections (20 μm thick) were cut on a cryostat (Leica CM1950) and mounted onto charged glass slides (Fisher, 22-037-246), then stored at –80°C until use.

#### Immunostaining procedure

Slides were brought to room temperature, and a hydrophobic barrier was drawn around each section using a Super PAP Pen. Sections were rehydrated in PBS for 10 minutes and blocked in 10% normal goat serum containing 0.3% Triton X-100 (Sigma, T9284) for 1 hour at room temperature.

Sections were incubated overnight at 4°C with the following primary antibodies diluted in 5% serum with 0.3% Triton X-100:

| Primary Antibody | Target | Dilution | Vendor / Cat. No. |
| --- | --- | --- | --- |
| His-tag (Alexa Fluor 488) | Nanobody localization | 1:250 | CST, 14930 |
| Anti-IBA1 (SPICA Dye 594) | Microglia | 1:250 | Wako, 012-28401 |
| Anti-GFAP | Astrocytes | 1:250 | BD Biosciences, 556327 |
| Anti-human TfR1 | Human TfR1 | 1:250 | CST, 54505S |

After three washes in PBS containing 0.3% Triton X-100, sections were incubated for 2 hours at room temperature with appropriate fluorescent secondary antibodies diluted 1:1000 in PBS with 5% serum and 0.3% Triton X-100. The following secondary antibody was used:

| Secondary Antibody | Target | Dilution | Vendor / Cat. No. |
| --- | --- | --- | --- |
| Goat anti-Mouse IgG (Alexa Fluor 594) | GFAP, TfR1 | 1:1000 | Invitrogen, A-11032 |

Sections were washed in PBS and mounted using DAPI-containing aqueous mounting medium (Southern Biotech, 0100-20) and coverslipped (Corning, 2980-225).

#### Imaging and analysis

Images were captured using a Leica Stellaris 8 tau STED confocal microscope. Z-stacks were acquired with identical settings across all experimental conditions and processed LAS X software.

## Statistical Analysis

All quantitative data are presented as mean ± standard error of the mean (SEM), unless otherwise indicated. Statistical analyses were performed using GraphPad Prism software. Comparisons between two groups were made using unpaired two-tailed Student’s t-tests. For multiple group comparisons, one-way or two-way ANOVA followed by appropriate post hoc tests (e.g., Tukey’s or Sidak’s) were used. A p-value < 0.05 was considered statistically significant. The number of biological replicates (n) is indicated in the figure legends or methods.

## Data and code availability

All data supporting the findings of this study are available in the Supporting Values Excel file accompanying this paper.

## Acknowledgments

This study was funded by the National Institute on Aging (NIA).

## Author contributions

L.D. conceived and supervised the study. L.D., T.Y., S.M., and M.Y. designed the experiments. Experiments were performed by L.D., T.Y., S.M., and M.Y. The manuscript was written by L.D. and T.Y.

## References

1. Fillit, H., Ding, W. H., Buee, L., Kalman, J., Altstiel, L., Lawlor, B. et al. (1991) Elevated circulating tumor necrosis factor levels in Alzheimer’s disease Neurosci Lett 129, 318–320 10.1016/0304-3940(91)90490-k

2. Tarkowski, E., Andreasen, N., Tarkowski, A., and Blennow, K. (2003) Intrathecal inflammation precedes development of Alzheimer’s disease J Neurol Neurosurg Psychiatry 74, 1200–1205 10.1136/jnnp.74.9.1200

3. Tarkowski, E., Liljeroth, A. M., Minthon, L., Tarkowski, A., Wallin, A., and Blennow, K. (2003) Cerebral pattern of pro- and anti-inflammatory cytokines in dementias Brain Res Bull 61, 255–260 10.1016/s0361-9230(03)00088-1

4. Tarkowski, E., Tullberg, M., Fredman, P., and Wikkelso, C. (2003) Correlation between intrathecal sulfatide and TNF-alpha levels in patients with vascular dementia Dement Geriatr Cogn Disord 15, 207–211 10.1159/000068780

5. Alvarez, A., Cacabelos, R., Sanpedro, C., García-Fantini, M., and Aleixandre, M. (2007) Serum TNF-alpha levels are increased and correlate negatively with free IGF-I in Alzheimer disease Neurobiol Aging 28, 533–536 10.1016/j.neurobiolaging.2006.02.012

6. Plantone, D., Pardini, M., Righi, D., Manco, C., Colombo, B. M., and De Stefano, N. (2023) The Role of TNF-alpha in Alzheimer’s Disease: A Narrative Review Cells 13, 10.3390/cells13010054

7. Stellwagen, D., and Malenka, R. C. (2006) Synaptic scaling mediated by glial TNF-alpha Nature 440, 1054–1059 10.1038/nature04671

8. Stellwagen, D., Beattie, E. C., Seo, J. Y., and Malenka, R. C. (2005) Differential regulation of AMPA receptor and GABA receptor trafficking by tumor necrosis factor-alpha J Neurosci 25, 3219–3228 10.1523/JNEUROSCI.4486-04.2005

9. Beattie, E. C., Stellwagen, D., Morishita, W., Bresnahan, J. C., Ha, B. K., Von Zastrow, M. et al. (2002) Control of synaptic strength by glial TNFalpha Science 295, 2282–2285 10.1126/science.1067859

10. Cirrito, J. R., Kang, J. E., Lee, J., Stewart, F. R., Verges, D. K., Silverio, L. M. et al. (2008) Endocytosis is required for synaptic activity-dependent release of amyloid-beta in vivo Neuron 58, 42–51 S0896-6273(08)00124-4 [pii] 10.1016/j.neuron.2008.02.003

11. Schroeder, B. E., and Koo, E. H. (2005) To think or not to think: synaptic activity and Abeta release Neuron 48, 873–875 10.1016/j.neuron.2005.12.005

12. Cirrito, J. R., Yamada, K. A., Finn, M. B., Sloviter, R. S., Bales, K. R., May, P. C. et al. (2005) Synaptic activity regulates interstitial fluid amyloid-beta levels in vivo Neuron 48, 913–922 10.1016/j.neuron.2005.10.028

13. Crimins, J. L., Pooler, A., Polydoro, M., Luebke, J. I., and Spires-Jones, T. L. (2013) The intersection of amyloid beta and tau in glutamatergic synaptic dysfunction and collapse in Alzheimer’s disease Ageing Res Rev 12, 757–763 10.1016/j.arr.2013.03.002

14. Pooler, A. M., Polydoro, M., Wegmann, S., Nicholls, S. B., Spires-Jones, T. L., and Hyman, B. T. (2013) Propagation of tau pathology in Alzheimer’s disease: identification of novel therapeutic targets Alzheimers Res Ther 5, 49 10.1186/alzrt214

15. Pooler, A. M., Polydoro, M., Maury, E. A., Nicholls, S. B., Reddy, S. M., Wegmann, S. et al. (2015) Amyloid accelerates tau propagation and toxicity in a model of early Alzheimer’s disease Acta Neuropathol Commun 3, 14 10.1186/s40478-015-0199-x

16. Pooler, A. M., Noble, W., and Hanger, D. P. (2014) A role for tau at the synapse in Alzheimer’s disease pathogenesis Neuropharmacology 76 Pt A, 1–8 10.1016/j.neuropharm.2013.09.018

17. Polydoro, M., Dzhala, V. I., Pooler, A. M., Nicholls, S. B., McKinney, A. P., Sanchez, L. et al. (2014) Soluble pathological tau in the entorhinal cortex leads to presynaptic deficits in an early Alzheimer’s disease model Acta Neuropathol 127, 257–270 10.1007/s00401-013-1215-5

18. Pooler, A. M., Phillips, E. C., Lau, D. H., Noble, W., and Hanger, D. P. (2013) Physiological release of endogenous tau is stimulated by neuronal activity EMBO Rep 14, 389–394 10.1038/embor.2013.15

19. Pooler, A. M., and Hanger, D. P. (2010) Functional implications of the association of tau with the plasma membrane Biochem Soc Trans 38, 1012–1015 10.1042/BST0381012

20. Akiyama, H., Barger, S., Barnum, S., Bradt, B., Bauer, J., Cole, G. M. et al. (2000) Inflammation and Alzheimer’s disease Neurobiol Aging 21, 383–421 10.1016/s0197-4580(00)00124-x

21. Guerreiro, R., Wojtas, A., Bras, J., Carrasquillo, M., Rogaeva, E., Majounie, E. et al. (2013) TREM2 variants in Alzheimer’s disease N Engl J Med 368, 117–127 10.1056/NEJMoa1211851

22. Jonsson, T., Stefansson, H., Steinberg, S., Jonsdottir, I., Jonsson, P. V., Snaedal, J. et al. (2013) Variant of TREM2 associated with the risk of Alzheimer’s disease N Engl J Med 368, 107–116 10.1056/NEJMoa1211103

23. Song, W. M., Joshita, S., Zhou, Y., Ulland, T. K., Gilfillan, S., and Colonna, M. (2018) Humanized TREM2 mice reveal microglia-intrinsic and -extrinsic effects of R47H polymorphism J Exp Med 215, 745–760 10.1084/jem.20171529

24. Tambini, M. D., and D’Adamio, L. (2020) Trem2 Splicing and Expression are Preserved in a Human Abeta-producing, Rat Knock-in Model of Trem2-R47H Alzheimer’s Risk Variant Sci Rep 10, 4122 10.1038/s41598-020-60800-1

25. Ren, S., Breuillaud, L., Yao, W., Yin, T., Norris, K. A., Zehntner, S. P. et al. (2021) TNF-alpha-mediated reduction in inhibitory neurotransmission precedes sporadic Alzheimer’s disease pathology in young Trem2(R47H) rats J Biol Chem 296, 100089 10.1074/jbc.RA120.016395

26. Ren, S., Yao, W., Tambini, M. D., Yin, T., Norris, K. A., and D’Adamio, L. (2020) Microglia TREM2(R47H) Alzheimer-linked variant enhances excitatory transmission and reduces LTP via increased TNF-alpha levels Elife 9, 10.7554/eLife.57513

27. Perry, R. T., Collins, J. S., Harrell, L. E., Acton, R. T., and Go, R. C. (2001) Investigation of association of 13 polymorphisms in eight genes in southeastern African American Alzheimer disease patients as compared to age-matched controls Am J Med Genet 105, 332–342 10.1002/ajmg.1371

28. Perry, R. T., Collins, J. S., Wiener, H., Acton, R., and Go, R. C. (2001) The role of TNF and its receptors in Alzheimer’s disease Neurobiol Aging 22, 873–883 10.1016/s0197-4580(01)00291-3

29. MacPherson, K. P., Sompol, P., Kannarkat, G. T., Chang, J., Sniffen, L., Wildner, M. E. et al. (2017) Peripheral administration of the soluble TNF inhibitor XPro1595 modifies brain immune cell profiles, decreases beta-amyloid plaque load, and rescues impaired long-term potentiation in 5xFAD mice Neurobiol Dis 102, 81–95 10.1016/j.nbd.2017.02.010

30. Montgomery, S. L., and Bowers, W. J. (2012) Tumor necrosis factor-alpha and the roles it plays in homeostatic and degenerative processes within the central nervous system J Neuroimmune Pharmacol 7, 42–59 10.1007/s11481-011-9287-2

31. Watad, A., McGonagle, D., Anis, S., Carmeli, R., Cohen, A. D., Tsur, A. M. et al. (2022) TNF inhibitors have a protective role in the risk of dementia in patients with ankylosing spondylitis: Results from a nationwide study Pharmacol Res 182, 106325 10.1016/j.phrs.2022.106325

32. Chou, R. C., Kane, M., Ghimire, S., Gautam, S., and Gui, J. (2016) Treatment for Rheumatoid Arthritis and Risk of Alzheimer’s Disease: A Nested Case-Control Analysis CNS Drugs 30, 1111–1120 10.1007/s40263-016-0374-z

33. Zhou, M., Xu, R., Kaelber, D. C., and Gurney, M. E. (2020) Tumor Necrosis Factor (TNF) blocking agents are associated with lower risk for Alzheimer’s disease in patients with rheumatoid arthritis and psoriasis PLoS One 15, e0229819 10.1371/journal.pone.0229819

34. Shi, J. Q., Wang, B. R., Jiang, W. W., Chen, J., Zhu, Y. W., Zhong, L. L. et al. (2011) Cognitive improvement with intrathecal administration of infliximab in a woman with Alzheimer’s disease J Am Geriatr Soc 59, 1142–1144 10.1111/j.1532-5415.2011.03445.x

35. Tobinick, E. (2012) Deciphering the physiology underlying the rapid clinical effects of perispinal etanercept in Alzheimer’s disease Curr Alzheimer Res 9, 99–109, http://www.ncbi.nlm.nih.gov/pubmed/22191562

36. Tobinick, E., Gross, H., Weinberger, A., and Cohen, H. (2006) TNF-alpha modulation for treatment of Alzheimer’s disease: a 6-month pilot study MedGenMed 8, 25, https://www.ncbi.nlm.nih.gov/pubmed/16926764

37. Tracey, D., Klareskog, L., Sasso, E. H., Salfeld, J. G., and Tak, P. P. (2008) Tumor necrosis factor antagonist mechanisms of action: a comprehensive review Pharmacol Ther 117, 244–279 10.1016/j.pharmthera.2007.10.001

38. Cheng, X., Shen, Y., and Li, R. (2014) Targeting TNF: a therapeutic strategy for Alzheimer’s disease Drug Discov Today 19, 1822–1827 10.1016/j.drudis.2014.06.029

39. Boado, R. J., Hui, E. K., Lu, J. Z., Zhou, Q. H., and Pardridge, W. M. (2010) Selective targeting of a TNFR decoy receptor pharmaceutical to the primate brain as a receptor-specific IgG fusion protein J Biotechnol 146, 84–91 10.1016/j.jbiotec.2010.01.011

40. (2015) Etanercept in Alzheimer disease: A randomized, placebo-controlled, double-blind, phase 2 trial Neurology 85, 2084 10.1212/WNL.0000000000002206

41. Yin, T., Ramon, A., Greenig, M., Sormanni, P., and D’Adamio, L. (2025) Development of potent humanized TNFalpha inhibitory nanobodies for therapeutic applications in TNFalpha-mediated diseases MAbs 17, 2498164 10.1080/19420862.2025.2498164

42. Yin, T., Yesiltepe, M., Metkar, S., Ramon, A., Greenig, M., Sormanni, P. et al. (2025) The NewroBus platform: engineered humanized anti-TfR1 nanobodies for efficient brain delivery Cell Commun Signal 24, 69 10.1186/s12964-025-02605-1

43. Johnsen, K. B., Burkhart, A., Thomsen, L. B., Andresen, T. L., and Moos, T. (2019) Targeting the transferrin receptor for brain drug delivery Prog Neurobiol 181, 101665 10.1016/j.pneurobio.2019.101665

44. Pardridge, W. M., Buciak, J. L., and Friden, P. M. (1991) Selective transport of an anti-transferrin receptor antibody through the blood-brain barrier in vivo J Pharmacol Exp Ther 259, 66–70, https://www.ncbi.nlm.nih.gov/pubmed/1920136

45. Lee, H. J., Engelhardt, B., Lesley, J., Bickel, U., and Pardridge, W. M. (2000) Targeting rat anti-mouse transferrin receptor monoclonal antibodies through blood-brain barrier in mouse J Pharmacol Exp Ther 292, 1048–1052, https://www.ncbi.nlm.nih.gov/pubmed/10688622

46. Ullman, J. C., Arguello, A., Getz, J. A., Bhalla, A., Mahon, C. S., Wang, J. et al. (2020) Brain delivery and activity of a lysosomal enzyme using a blood-brain barrier transport vehicle in mice Sci Transl Med 12, 10.1126/scitranslmed.aay1163

47. Weber, F., Bohrmann, B., Niewoehner, J., Fischer, J. A. A., Rueger, P., Tiefenthaler, G. et al. (2018) Brain Shuttle Antibody for Alzheimer’s Disease with Attenuated Peripheral Effector Function due to an Inverted Binding Mode Cell Rep 22, 149–162 10.1016/j.celrep.2017.12.019

48. Huang, Q., Chan, K. Y., Wu, J., Botticello-Romero, N. R., Zheng, Q., Lou, S. et al. (2024) An AAV capsid reprogrammed to bind human transferrin receptor mediates brain-wide gene delivery Science 384, 1220–1227 10.1126/science.adm8386

49. Yesiltepe, M., Metkar, S., Yin, T., Chakraborty, I., and D’Adamio, L. (2026) A humanized transferrin receptor 1-transferrin model supports functional iron homeostasis and therapeutic delivery across the blood-brain barrier J Biol Chem 302, 110995 10.1016/j.jbc.2025.110995

50. Kariolis, M. S., Wells, R. C., Getz, J. A., Kwan, W., Mahon, C. S., Tong, R. et al. (2020) Brain delivery of therapeutic proteins using an Fc fragment blood-brain barrier transport vehicle in mice and monkeys Sci Transl Med 12, 10.1126/scitranslmed.aay1359

51. Trenor, C. C., 3rd, Campagna, D. R., Sellers, V. M., Andrews, N. C., and Fleming, M. D. (2000) The molecular defect in hypotransferrinemic mice Blood 96, 1113–1118, https://www.ncbi.nlm.nih.gov/pubmed/10910930

52. Zalevsky, J., Secher, T., Ezhevsky, S. A., Janot, L., Steed, P. M., O’Brien, C. et al. (2007) Dominant-negative inhibitors of soluble TNF attenuate experimental arthritis without suppressing innate immunity to infection J Immunol 179, 1872–1883 10.4049/jimmunol.179.3.1872

53. Liddelow, S. A., Guttenplan, K. A., Clarke, L. E., Bennett, F. C., Bohlen, C. J., Schirmer, L. et al. (2017) Neurotoxic reactive astrocytes are induced by activated microglia Nature 541, 481–487 10.1038/nature21029

54. Liddelow, S. A., and Barres, B. A. (2017) Reactive Astrocytes: Production, Function, and Therapeutic Potential Immunity 46, 957–967 10.1016/j.immuni.2017.06.006

55. Fu, W., Hettinghouse, A., Chen, Y., Hu, W., Ding, X., Chen, M. et al. (2021) 14-3-3 epsilon is an intracellular component of TNFR2 receptor complex and its activation protects against osteoarthritis Ann Rheum Dis 80, 1615–1627 10.1136/annrheumdis-2021-220000

56. Zhang, N., Wang, Z., and Zhao, Y. (2020) Selective inhibition of Tumor necrosis factor receptor-1 (TNFR1) for the treatment of autoimmune diseases Cytokine Growth Factor Rev 55, 80–85 10.1016/j.cytogfr.2020.03.002

57. Hennessy, E., Griffin, E. W., and Cunningham, C. (2015) Astrocytes Are Primed by Chronic Neurodegeneration to Produce Exaggerated Chemokine and Cell Infiltration Responses to Acute Stimulation with the Cytokines IL-1beta and TNF-alpha J Neurosci 35, 8411–8422 10.1523/JNEUROSCI.2745-14.2015

58. Bezzi, P., Domercq, M., Brambilla, L., Galli, R., Schols, D., De Clercq, E. et al. (2001) CXCR4-activated astrocyte glutamate release via TNFalpha: amplification by microglia triggers neurotoxicity Nat Neurosci 4, 702–710 10.1038/89490

59. Fine, S. M., Angel, R. A., Perry, S. W., Epstein, L. G., Rothstein, J. D., Dewhurst, S. et al. (1996) Tumor necrosis factor alpha inhibits glutamate uptake by primary human astrocytes. Implications for pathogenesis of HIV-1 dementia J Biol Chem 271, 15303–15306 10.1074/jbc.271.26.15303

60. Hu, S., Sheng, W. S., Ehrlich, L. C., Peterson, P. K., and Chao, C. C. (2000) Cytokine effects on glutamate uptake by human astrocytes Neuroimmunomodulation 7, 153–159 10.1159/000026433

61. Tilleux, S., and Hermans, E. (2007) Neuroinflammation and regulation of glial glutamate uptake in neurological disorders J Neurosci Res 85, 2059–2070 10.1002/jnr.21325

62. Zou, J. Y., and Crews, F. T. (2005) TNF alpha potentiates glutamate neurotoxicity by inhibiting glutamate uptake in organotypic brain slice cultures: neuroprotection by NF kappa B inhibition Brain Res 1034, 11–24 10.1016/j.brainres.2004.11.014

63. Kim, H., Leng, K., Park, J., Sorets, A. G., Kim, S., Shostak, A. et al. (2022) Reactive astrocytes transduce inflammation in a blood-brain barrier model through a TNF-STAT3 signaling axis and secretion of alpha 1-antichymotrypsin Nat Commun 13, 6581 10.1038/s41467-022-34412-4

