## Supplementary material for "NN-800s: Brain-penetrant trimeric nanobodies enable potent TNFα inhibition via TfR1-mediated transcytosis": Suppl.

### Supplemental Material

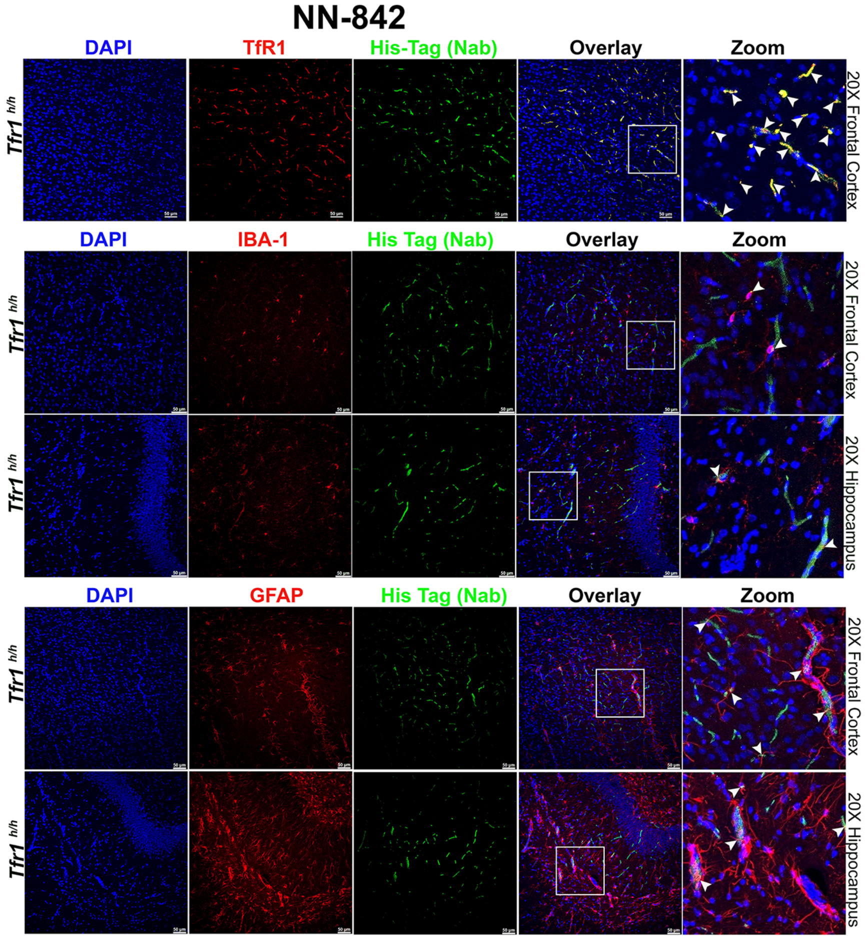


**Figure S1. Cellular localization of NN-842 in cortex and hippocampus.** Representative immunofluorescence of NN-842 in *Tfr1*$^{h/h}$ rat brain. NN-842 (green, anti-His), hTfR1 (red), and DAPI show signal in vascular and parenchymal compartments; arrows indicate colocalization (top). Brain sections (cortex and hippocampus) stained with IBA1 (microglia) or GFAP (astrocytes) demonstrate NN-842 localization in both cell types (middle and bottom). White arrows in the zoomed-in panel indicate the colocalization yellow spots within the region highlighted by the white square in the overlay panel.

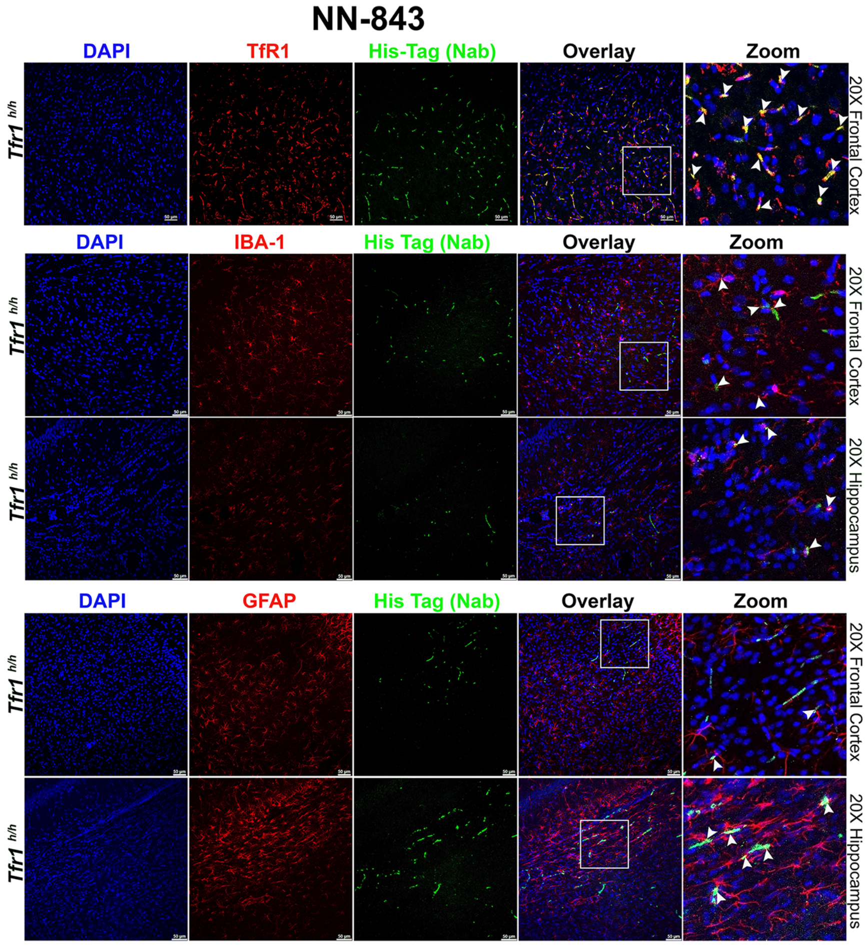


**Figure S2. Cellular localization of NN-843 in cortex and hippocampus.** Representative immunofluorescence of NN-843 in *Tfr1*$^{h/h}$ rat brain. NN-843 (green, anti-His), hTfR1 (red), and DAPI show signal in vascular and parenchymal compartments; arrows indicate colocalization (top). Brain sections (cortex and hippocampus) stained with IBA1 (microglia) or GFAP (astrocytes) demonstrate NN-843 localization in both cell types (middle and bottom). White arrows in the zoomed-in panel indicate the colocalization yellow spots within the region highlighted by the white square in the overlay panel.

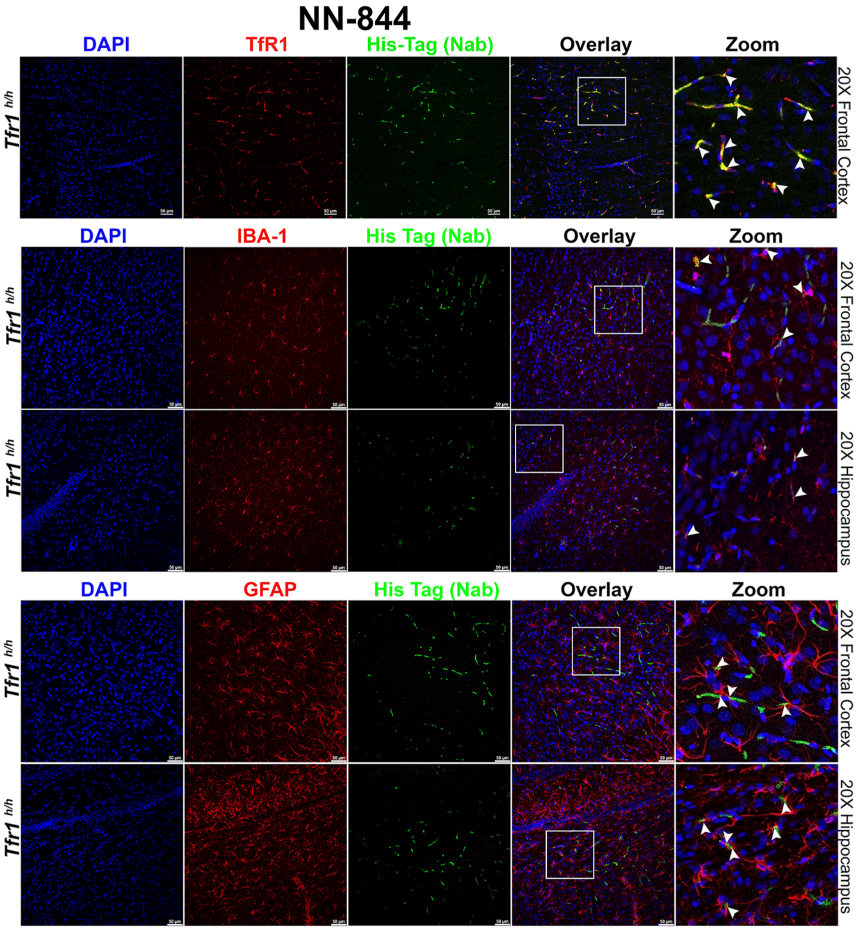


**Figure S3. Cellular localization of NN-843 in cortex and hippocampus.** Representative immunofluorescence of NN-844 in *Tfr1*$^{h/h}$ rat brain. NN-844 (green, anti-His), hTfR1 (red), and DAPI show signal in vascular and parenchymal compartments; arrows indicate colocalization (top). Brain sections (cortex and hippocampus) stained with IBA1 (microglia) or GFAP (astrocytes) demonstrate NN-844 localization in both cell types (middle and bottom). White arrows in the zoomed-in panel indicate the colocalization yellow spots within the region highlighted by the white square in the overlay panel.

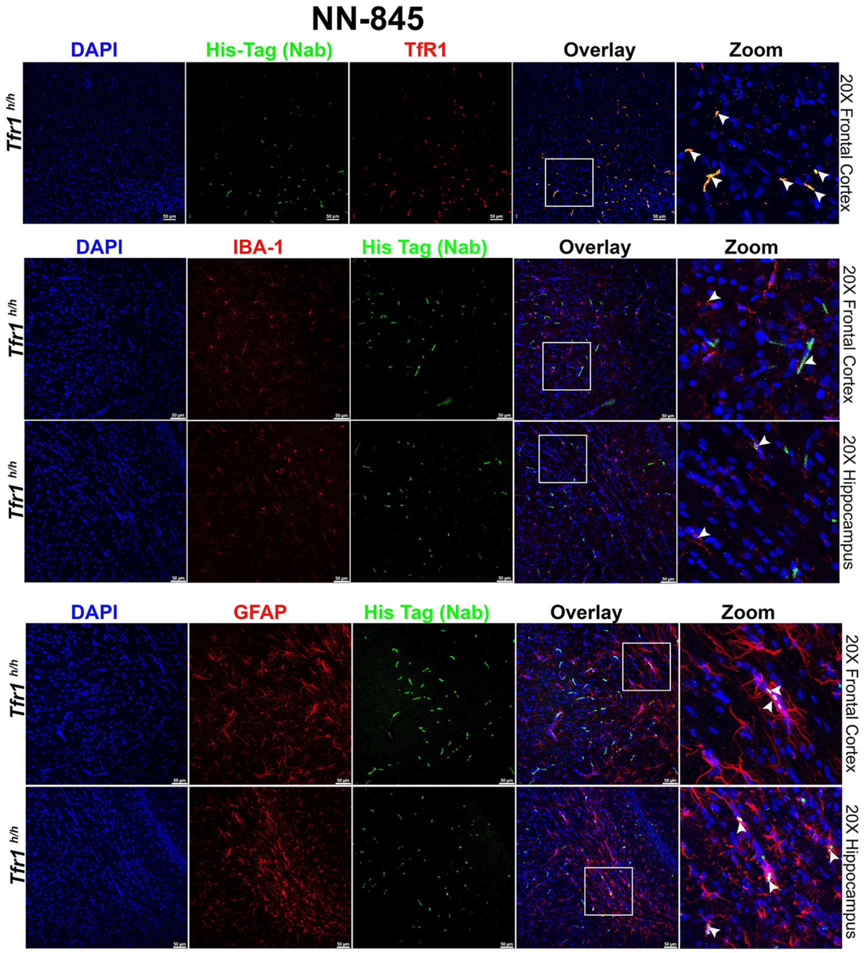


**Figure S4. Cellular localization of NN-845 in cortex and hippocampus.** Representative immunofluorescence of NN-845 in *Tfr1*$^{h/h}$ rat brain. NN-845 (green, anti-His), hTfR1 (red), and DAPI show signal in vascular and parenchymal compartments; arrows indicate colocalization (top). Brain sections (cortex and hippocampus) stained with IBA1 (microglia) or GFAP (astrocytes) demonstrate NN-845 localization in both cell types (middle and bottom). White arrows in the zoomed-in panel indicate the colocalization yellow spots within the region highlighted by the white square in the overlay panel.

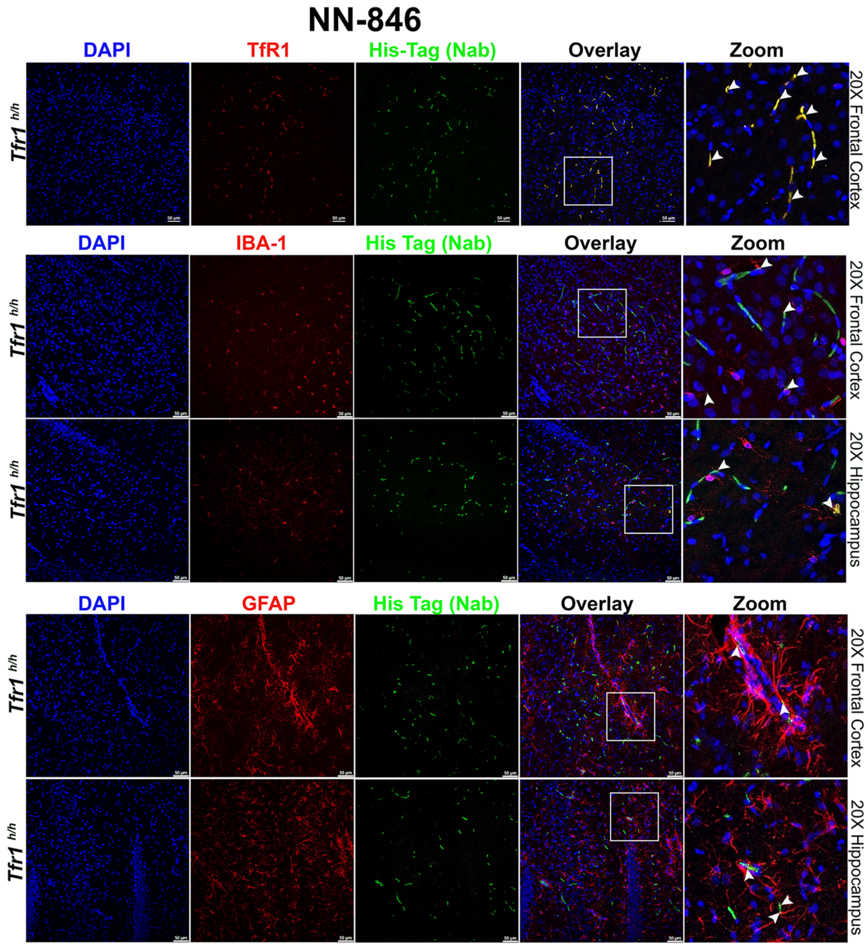


**Figure S5. Cellular localization of NN-846 in cortex and hippocampus.** Representative immunofluorescence of NN-846 in *Tfr1*$^{h/h}$ rat brain. NN-846 (green, anti-His), hTfR1 (red), and DAPI show signal in vascular and parenchymal compartments; arrows indicate colocalization (top). Brain sections (cortex and hippocampus) stained with IBA1 (microglia) or GFAP (astrocytes) demonstrate NN-846 localization in both cell types (middle and bottom). White arrows in the zoomed-in panel indicate the colocalization yellow spots within the region highlighted by the white square in the overlay panel.

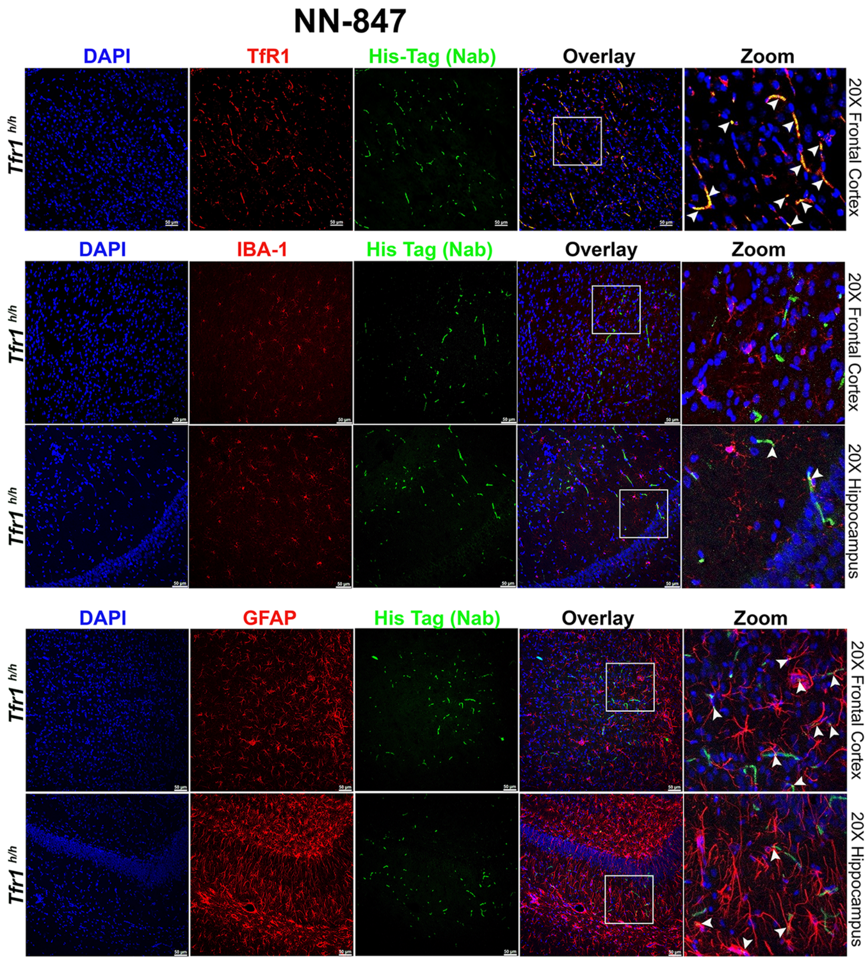


**Figure S6. Cellular localization of NN-847 in cortex and hippocampus.** Representative immunofluorescence of NN-847 in *Tfr1*$^{h/h}$ rat brain. NN-847 (green, anti-His), hTfR1 (red), and DAPI show signal in vascular and parenchymal compartments; arrows indicate colocalization (top). Brain sections (cortex and hippocampus) stained with IBA1 (microglia) or GFAP (astrocytes) demonstrate NN-847 localization in both cell types (middle and bottom). White arrows in the zoomed-in panel indicate the colocalization yellow spots within the region highlighted by the white square in the overlay panel.

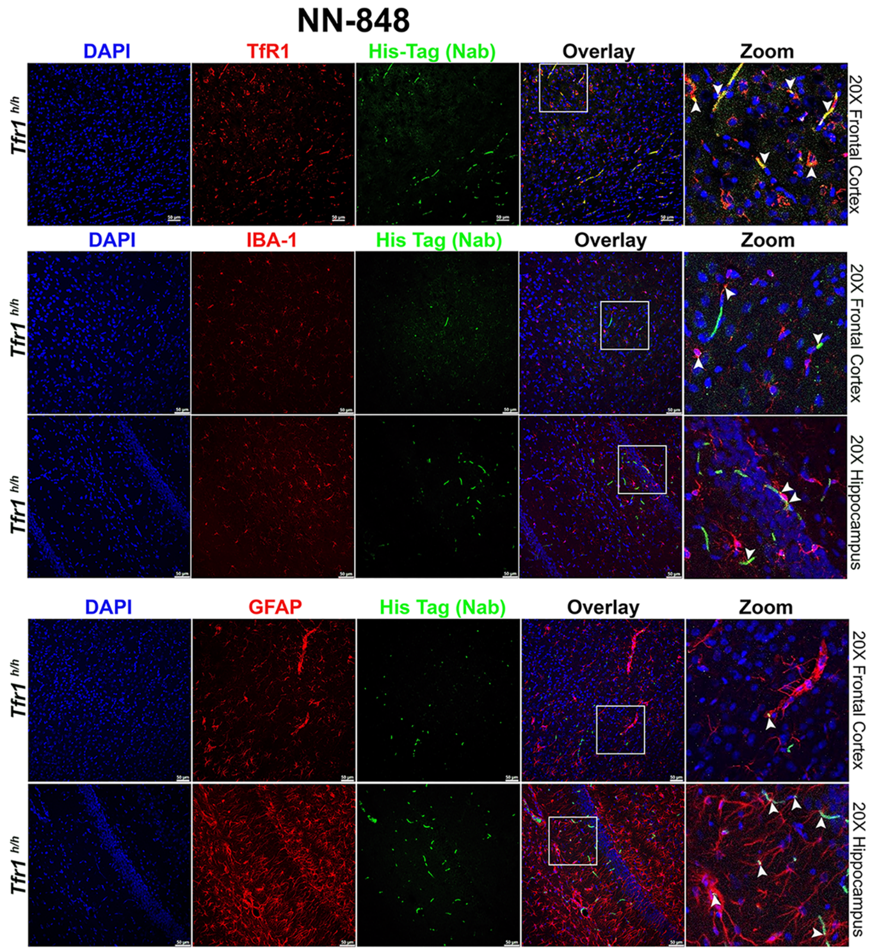


**Figure S7. Cellular localization of NN-848 in cortex and hippocampus.** Representative immunofluorescence of NN-848 in *Tfr1*$^{h/h}$ rat brain. NN-848 (green, anti-His), hTfR1 (red), and DAPI show signal in vascular and parenchymal compartments; arrows indicate colocalization (top). Brain sections (cortex and hippocampus) stained with IBA1 (microglia) or GFAP (astrocytes) demonstrate NN-848 localization in both cell types (middle and bottom). White arrows in the zoomed-in panel indicate the colocalization yellow spots within the region highlighted by the white square in the overlay panel.
